# Single-component, self-assembling, protein nanoparticles presenting the receptor binding domain and stabilized spike as SARS-CoV-2 vaccine candidates

**DOI:** 10.1101/2020.09.14.296715

**Authors:** Linling He, Xiaohe Lin, Ying Wang, Ciril Abraham, Cindy Sou, Timothy Ngo, Yi Zhang, Ian A. Wilson, Jiang Zhu

## Abstract

Vaccination against SARS-CoV-2 provides an effective tool to combat the COIVD-19 pandemic. Here, we combined antigen optimization and nanoparticle display to develop vaccine candidates for SARS-CoV-2. We first displayed the receptor-binding domain (RBD) on three self-assembling protein nanoparticle (SApNP) platforms using the SpyTag/SpyCatcher system. We then identified heptad repeat 2 (HR2) in S2 as the cause of spike metastability, designed an HR2-deleted glycine-capped spike (S2GΔHR2), and displayed S2GΔHR2 on SApNPs. An antibody column specific for the RBD enabled tag-free vaccine purification. In mice, the 24-meric RBD-ferritin SApNP elicited a more potent neutralizing antibody (NAb) response than the RBD alone and the spike with two stabilizing proline mutations in S2 (S2P). S2GΔHR2 elicited two-fold-higher NAb titers than S2P, while S2GΔHR2 SApNPs derived from multilayered E2p and I3-01v9 60-mers elicited up to 10-fold higher NAb titers. The S2GΔHR2-presenting I3-01v9 SApNP also induced critically needed T-cell immunity, thereby providing a promising vaccine candidate.

**ONE-SENTENCE SUMMARY:** The SARS-CoV-2 receptor binding domain and S2GΔHR2 spike elicited potent immune responses when displayed on protein nanoparticles as vaccine candidates.

## INTRODUCTION

Three coronaviruses (CoVs) have caused widespread outbreaks in humans, including severe acute respiratory syndrome CoV-1 (SARS-CoV-1), Middle East respiratory syndrome CoV (MERS-CoV), and SARS-CoV-2, which is the causative agent of COVID-19 (*1–3*) and has resulted in more than 1.9 million deaths worldwide (*4*). Enormous efforts are being undertaken to develop effective therapeutics and prophylactics for SARS-CoV-2. Small molecules that can block the host receptor, angiotensin-converting enzyme 2 (ACE2), and the transmembrane protease serine 2 (TMPRSS2) (*5*), which is required to process the spike protein, are being considered as treatments in addition to other interventions (*6*). While the immunology underlying COVID-19 is still being intensively studied (*6–8*), various vaccine candidates are now in clinical development (*9–12*). Inactivated virus vaccines have exhibited robust neutralizing antibody (NAb) responses in animals (*13, 14*), whereas viral vector vaccines based on human adenovirus (Ad5 and Ad26) and chimpanzee ChAdOx1 have been evaluated in nonhuman primates (NHPs) and human trials (*15–18*). Both DNA (*19–21*) and mRNA (*22, 23*) vaccines have been rapidly developed, with moderate NAb titers observed for the mRNA vaccine in medium and high dose groups (*22*). A recombinant spike protein adjuvanted with lipid nanoparticles (NPs), NVX-CoV2373, elicited high NAb titers in a human trial that were on average four-fold greater than in convalescent patients (*24, 25*). Efficacy was recently reported for a vector vaccine (AZD1222: 70.4%) (*26*) and two mRNA vaccines (mRNA-1273: 94.1% and BNT162b2: 95%) (*27, 28*). In December 2020, the U.S. Food and Drug Administration (FDA) issued the emergency use authorization (EUA) for the two mRNA vaccines. The SARS-CoV-2 spike protein is a trimer of S1-S2 heterodimers. The S1 subunit contains a receptor-binding domain (RBD) that binds to ACE2 on host cells to initiate infection.

The S2 subunit consists of a fusion peptide (FP) and heptad repeat regions 1 and 2 (HR1 and HR2). Upon endocytosis of the virion, the S1 subunit is cleaved off to facilitate FP insertion into the host cell membrane, while the remaining S2 refolds to bring HR1 and HR2 together to fuse the viral and host cell membranes (*29*). The spike protein harbors all NAb epitopes and is the main target for vaccine development against SARS-associated CoVs (*30*). Convalescent plasma (CP) has been used to treat COVID-19 patients with severe conditions (*31*), highlighting the importance of NAbs in protection (*32*). Due to moderate sequence conservation of the RBDs (∼73%), some previously identified NAbs targeting the SARS-CoV-1 RBD have been shown to bind and cross-neutralize SARS-CoV-2 (*33, 34*). Using single-cell technologies and the SARS-CoV-2 RBD or spike as bait, potent NAbs have now been isolated from COVID-19 patients (*35–41*). Camelid-derived single-chain NAbs have also been obtained by panning naïve or immune llama single-chain antibody (VHH) libraries (*42, 43*). Structures of the SARS-CoV-2 spike and RBD in unliganded (*44, 45*), ACE2-bound (*46–48*), and antibody-bound (*49–51*) states determined by x-ray crystallography and cryo-electron microscopy (cryo-EM) have paved the way for rational vaccine design. Cryo-EM and cryo-electron tomography (ET) have revealed the inherent spike metastability and the co-existence of pre/post-fusion spikes on virions (*52*). A double-proline mutation (*52*) has been used in most soluble spike (S2P) constructs and all but inactivated vaccines, although a HexaPro version with greater yield and stability is now available (*53*). Cryo-ET has also uncovered a dynamic, triple-hinged HR2 stalk that facilitates viral entry and immune evasion (*54–56*).

In this study, we designed and optimized SARS-CoV-2 antigens for multivalent display on self-assembling protein nanoparticles (SApNPs) (*57–59*), including a ferritin (FR) 24-mer and two 60-mers (E2p and I3-01) containing an inner layer of locking domains (LD) and a cluster of T-cell epitopes (*60*). To facilitate tag-free vaccine purification, we developed an immunoaffinity column based on antibody CR3022 that binds to both SARS-CoV-1/2 RBDs (*34, 50*). We first designed a scaffolded RBD trimer construct to mimic the “RBD-up” spike conformation. The SARS-CoV-1/2 RBDs were attached to SApNPs using the SpyTag/SpyCatcher system (*61*), providing a robust strategy for developing RBD-based nanoparticle vaccines. We then probed the spike metastability by comparing two uncleaved spike antigens, S2P (K986P/V987P) and S2G (K986G/V987G). The SARS-CoV-2 S2G spike exhibited abnormal behavior, suggesting that an unidentified facet of the spike can promote conformational change and block antibody access to the RBD. An HR2-deleted spike, S2GΔHR2, produced high-purity trimers, suggesting that the HR2 stalk may be a trigger of spike metastability consistent with recent findings (*54–56*). We next displayed S2GΔHR2 on three SApNPs by gene fusion, resulting in spike SApNPs with high yield, purity, and antigenicity. In mouse immunization, the S2P spike protein elicited the lowest level of NAb response. In contrast, the scaffolded RBD trimer registered two-to-three-fold higher NAb titers, with another five-fold increase in NAb titer achieved by multivalent display on FR. S2GΔHR2 elicited up to seven-fold higher NAb titers, while the large, multilayered S2GΔHR2 E2p and I3-01 SApNPs induced up-to-10-fold higher NAb titers than S2P. Further analysis indicated that the S2GΔHR2-presenting I3-01v9 SApNP can elicit a strong Th1 response as well as other types of T-cell responses needed for protective cellular immunity. Our study thus identifies the HR2 stalk as a major source of spike metastability, validates an HR2-deleted spike design, and provides a set of RBD- and spike-based virus-like particles (VLPs) as effective protein vaccine candidates against SARS-CoV-2.

## RESULTS

### Rational design of scaffolded RBD trimers and RBD-presenting SApNPs

RBD binding to the ACE2 receptor initiates the membrane fusion process (*5*). The crystal structure of SARS-CoV-2 RBD/ACE2 complex revealed the atomic details of receptor recognition (*62*). The SARS-CoV-2 RBD has been used as bait to isolate monoclonal antibodies (mAbs) from patient samples (*35–41*). For SARS-CoV-1 and MERS-CoV, RBD-based vaccines have induced potent NAbs that effectively block viral entry (*30*). Therefore, the RBD represents a major target for humoral responses following infection and can be used to develop epitope-focused vaccines.

We first hypothesized that RBD attached to a trimeric scaffold could mimic the “RBD-up” spike conformation and elicit NAbs that block ACE2 binding. To test this possibility, we designed a fusion construct containing SARS-CoV-1/2 RBD, a short 5-aa G_4_S linker (with a 2-aa restriction site), and a trimeric viral capsid protein, SHP (PDB: 1TD0) (**Fig. 1A**). Structural modeling showed that the three tethered RBDs form a triangle of 92 Å (measured at L492), which is 14 and 18 Å wider than the SARS-CoV-1 “two-RBD-up” spike (PDB: 6CRX, measured at L478) (*63*) and the MERS-CoV “all-RBD-up” spike (PDB: 5X59, measured for L506) (*64*), respectively, allowing NAb access to each RBD. We then developed an immunoaffinity chromatography (IAC) column to facilitate tag-free purification. Previously, NAb-derived IAC columns have been used to purify HIV-1 Env trimers/NPs (*58, 59, 65, 66*), hepatitis C virus (HCV) E2 cores/NPs (*57*), and Ebola virus (EBOV) GP trimers/NPs (*60*). Tian et al. reported that a SARS-CoV-1 NAb, CR3022, can bind SARS-CoV-2 RBD (*34*). The SARS-CoV-2 RBD/CR3022 structure revealed a conserved cryptic epitope that is shared by the two SARS-CoVs, suggesting that transient breathing motions of the spike protein enabled CR3022 binding to the RBD (*50*). Here, we examined the utility of CR3022 in IAC columns. The SARS-CoV-1/2 RBD-5GS-1TD0 constructs were transiently expressed in 100-ml ExpiCHO cells and purified on a CR3022 column prior to size-exclusion chromatography (SEC) using a Superdex 200 10/300 GL column. While the SARS-CoV-1 RBD construct showed both aggregate (∼8.6 ml) and trimer (∼12.7 ml) peaks in the SEC profile, the SARS-CoV-2 RBD construct produced a single, pure trimer peak at ∼12.8 ml (**Fig. 1B**). In sodium dodecyl sulfate-polyacrylamide gel electrophoresis (SDS-PAGE), a monomer band of ∼37 kD and a trimer band of ∼100 kD were observed under reducing and non-reducing conditions, respectively (**fig. S1A**). Antigenicity was assessed for the two scaffolded RBD trimers in enzyme-linked immunosorbent assay (ELISA) after CR3022/SEC purification (**Fig. 1C**). RBD-specific NAbs targeting SARS-CoV-1 (CR3022 (*67*), m396 (*68*), 80R (*69*), and S230 (*70*)) and SARS-CoV-2 (B38 (*38*), CB6 (*37*), S309 from a SARS survivor (*33*), and P2B-2F6 (*36*)), were tested in ELISA. Overall, similar half maximal effective concentration (EC_50_) values were observed for the two RBD trimers binding to their respective NAbs (**Fig. 1C**). The SARS-CoV-1 RBD trimer showed greater affinity for CR3022 than its SARS-CoV-2 counterpart with a 1.3-fold difference in EC_50_ values, consistent with previous findings (*34, 50*). Of the SARS-CoV-2 NAbs, B38 yielded a similar EC_50_ value to CR3022, which can bind but not neutralize SARS-CoV-2. Antibody binding kinetics was measured by biolayer interferometry (BLI) (**Fig. 1D** and **Fig. 1B**). Overall, all tested mAbs exhibited a fast on-rate, but with visible differences in their off-rates. B38 showed a faster off-rate than other SARS-CoV-2 NAbs, while CR3022, the mAb used to purify SARS-CoV-1/2 RBD proteins, exhibited a comparable kinetic profile.

**Fig. 1.**
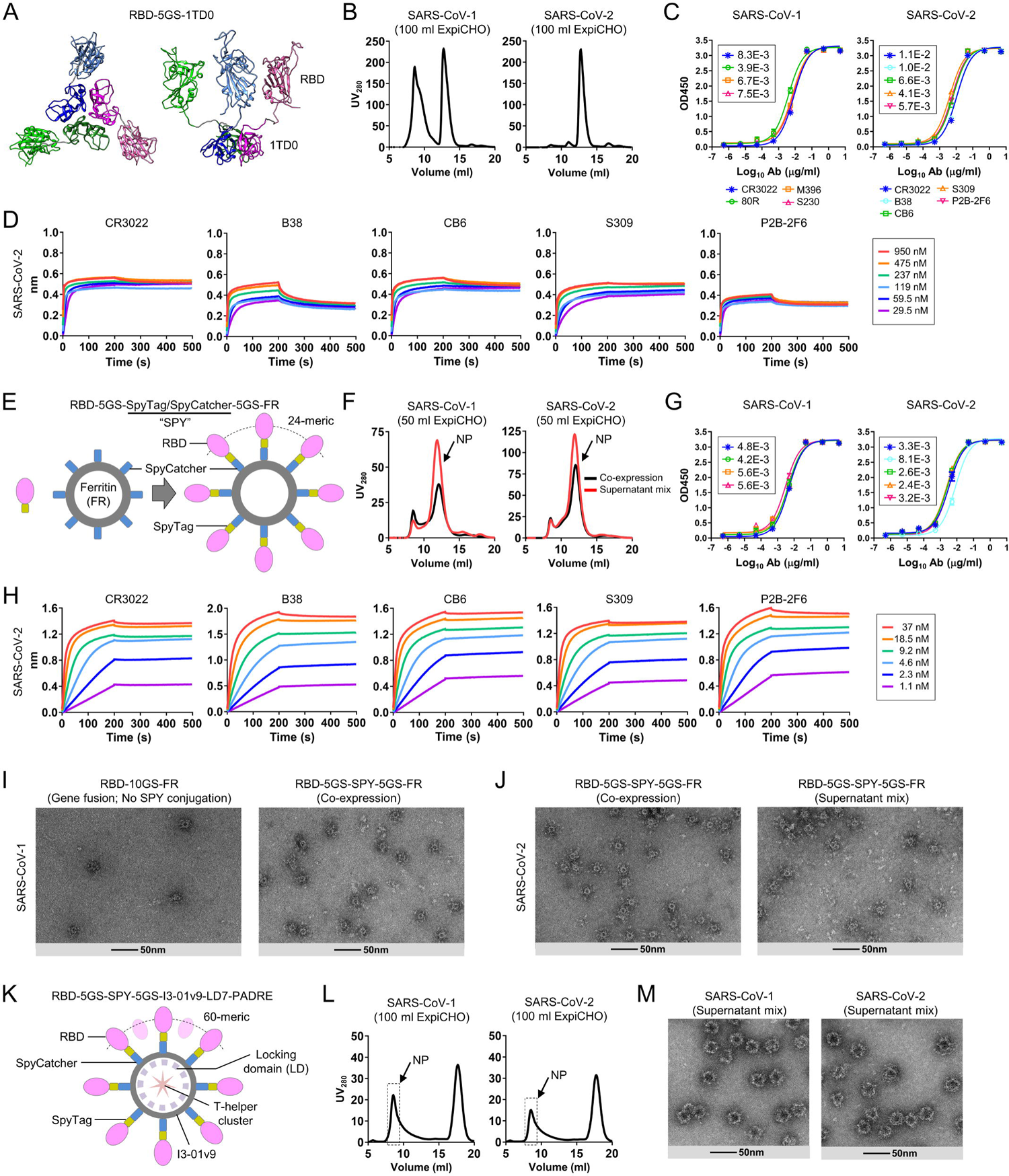
Rational design of SARS-CoV-1/2 RBD-based vaccines. **(A)** Structural model of RBD-5GS-1TD0 in an extended RBD-up conformation (top view and side view). 1TD0 is a trimerization scaffold of viral origin. **(B)** SEC profiles of SARS-CoV-1/2 scaffolded RBD trimers following ExpiCHO expression and CR3022 purification. **(C)** ELISA binding of SARS-CoV-1/2 scaffolded RBD trimers to a panel of mAbs. EC_50_ (μg/ml) values are labeled. **(D)** Octet binding of the SARS-CoV-2 scaffolded RBD trimer to five mAbs. Sensorgrams were obtained from an Octet RED96 instrument at six antigen concentrations from 950 to 29.5 nM by twofold dilutions. **(E)** Diagram of conjugating RBD to the 24-meric FR SApNP using the SpyTag/SpyCatcher (SPY) system. **(F)** SEC profiles of SARS-CoV-1/2 RBD-5GS-SPY-5GS-FR SApNPs produced in ExpiCHO by co-expression (black line) and supernatant mix (red line). **(G)** ELISA binding of SARS-CoV-1/2 RBD-FR SApNPs to a panel of mAbs. EC_50_ (μg/ml) values are labeled. **(H)** Octet binding of the SARS-CoV-2 RBD-FR SApNP to five mAbs. Sensorgrams were obtained from an Octet RED96 instrument at six antigen concentrations from 37 to 1.1 nM by twofold dilutions. **(I)** EM images of SARS-CoV-1 RBD-10GS-FR (gene fusion) and RBD-5GS-SPY-5GS-FR (co-expression). **(J)** EM images of SARS-CoV-2 RBD-5GS-SPY-5GS-FR produced by co-expression and supernatant mix. **(K)** Diagram of conjugating RBD to the 60-meric multilayered I3-01v9 SApNP using the SPY system. Locking domains (LD) and T-helper epitopes within the multilayered SApNP are depicted. **(L)** and **(M)** SEC profiles and EM images of SARS-CoV-1/2 RBD-5GS-SPY-5GS-I3-01v9-LD7-PADRE (or I3-01v9-L7P) SApNPs produced by supernatant mix. SEC profiles of scaffolded RBD trimers in (B) and SPY-linked RBD SApNPs in (F) and (L) were obtained from a Superdex 200 10/300 GL column and a Superose 6 10/300 GL column, respectively.

We then hypothesized that the SpyTag/SpyCatcher (termed SPY) system could be used to conjugate RBD to SApNPs to create multivalent RBD vaccines capable of eliciting a more potent NAb response (**Fig. 1E**). The 13-aa SpyTag spontaneously reacts with the SpyCatcher protein to form an irreversible isopeptide bond (*61*). The SPY system has been successfully used to attach antigens to VLPs (*71*). Here, SpyTag was fused to the C terminus of RBD, while SpyCatcher was fused to the N terminus of an SApNP subunit, both with a 5-aa G_4_S linker. This design was first tested for the 24-meric ferritin (FR). Here, we compared two production strategies, co-expression of RBD-5GS-SpyTag and SpyCatcher-5GS-FR versus supernatant mix after separate expression, both followed by purification on a CR3022 column. Protein obtained from transient transfection in 50-ml ExpiCHO cells was analyzed by SEC on a Superose 6 10/300 GL column (**Fig. 1F**). Both production strategies produced a peak (12 ml) corresponding to SApNPs. While the SARS-CoV-2 construct outperformed its SARS-CoV-1 counterpart in particle yield (0.6-1.0 mg versus 0.3-0.5 mg after CR3022/SEC), the supernatant mix appeared to be superior to co-expression for yield in both cases. The results thus suggest that both strategies can be used to produce RBD SApNPs in Chinese hamster ovary (CHO) cells in an industrial setting, as this mammalian expression system has been widely used to manufacture therapeutic glycoproteins under good manufacturing practice (GMP) conditions (*72, 73*). Antigenicity was assessed for SEC-purified RBD-5GS-SPY-5GS-FR SApNPs. In ELISA, RBD SApNPs showed slightly improved mAb binding compared to the RBD trimers, as indicated by EC_50_ values (**Fig. 1G**). In BLI, a more pronounced effect of multivalent display on antigenicity was observed, showing notably increased binding signals and plateaued dissociation (**Fig. 1H** and **Fig. 1C**). Structural integrity of various RBD SApNPs was analyzed by negative stain EM (nsEM) (**Figs. 1I** and **1J**). For SARS-CoV-1, a genetically fused RBD-10GS-FR construct produced very few, albeit highly pure, SApNPs (**Fig. 1I**, left). In contrast, the RBD-5GS-SPY-5GS-FR construct produced a high yield of SApNPs with visible surface decorations (**Fig. 1I**, right). For SARS-CoV-2, the purified RBD-5GS-SPY-5GS-FR SApNPs, irrespective of the production strategy, showed morphology of well-formed particles (**Fig. 1J**). Previously, we genetically fused a small LD protein and a T-cell epitope to each subunit of the E2p and I3-01v9 60-mers, resulting in “multilayered” SApNPs (*60*). PADRE, a 13-aa pan-DR epitope that activates CD4^+^ T cells (*74*), was used to promote B cell development toward NAbs. Here, two of the best designs, E2p-LD4-PADRE (or E2p-L4P) and I3-01v9-LD7-PADRE (or I3-01v9-L7P), were tested for their ability to display SARS-CoV-1/2 RBDs. Following the strategy established for FR, SARS-CoV-1/2 RBDs were attached to the I3-01v9-L7P SApNP using the SPY system (**Fig. 1K**). Despite the modest yield in ExpiCHO cells (**Fig. 1L**), large and pure particles were observed in the EM images (**Fig. 1M**). However, some impurities were noted for the RBD-presenting E2p-L4P SApNPs (**fig. S1D**). Nonetheless, we illustrated the utility of the SPY system for rapid development of SARS-CoV-1/2 RBD vaccines based on three different SApNP platforms.

Recently, RBDs were displayed on various SApNPs using the SPY system as MERS-CoV and SARS-CoV-2 vaccine candidates (*75, 76*). Walls et al. also reported a SARS-CoV-2 RBD vaccine candidate based on the two-component NP platform (*77*), which requires a “connector” component to facilitate NP assembly (*78*), likely resulting in suboptimal stability. Additionally, all these vaccine candidates require a complex production process involving two expression systems, separate purification steps, and an *in vitro* assembly step before final purification. In contrast, our SPY-linked RBD SApNPs are single-component by nature and can be produced in CHO cells with a simple purification scheme, offering unique advantages in stability and manufacturability.

### Rational design of prefusion spike through minimizing metastability

In addition to the RBD, the SARS-CoV-1/2 spikes contain other NAb epitopes (*30*), which are all presented in a trimeric context (**Fig. 2A**). A double-proline mutation (2P) between HR1 and the central helix (CH) has been used to stabilize the MERS-CoV (*79*) and SARS-CoV-1 spikes (*63*). A similar 2P mutation (K986P/V987P) was introduced into the SARS-CoV-2 spike (termed S2P), which has been used to isolate and characterize NAbs (*33, 35, 40, 42-45, 49*) and is the antigen in almost all vaccine candidates in clinical development (*11, 12*). However, a recent cryo-EM study revealed an unexpected packing of S1 in the S2P spike, positioned ∼12 Å outwards, compared to the full-length native spike, as well as a more ordered FP proximal region (FPPR) in S2 (*52*). New designs have been generated to control the spike conformation (*80*) or to further stabilize it with more prolines (HexaPro) (*53*). Recent cryo-EM and cryo-ET studies revealed that the SARS-CoV-2 spikes could adopt diverse orientations on native virions due to the highly flexible HR2 stalk (*54–56*). Previously, we identified an HR1 bend as the cause of HIV-1 Env metastability (*58, 81*) and examined the role of an equivalent HR1 bend and the HR2 stalk in EBOV GP metastability (*60*). This understanding of metastability proved critical for designing stable trimers and trimer-presenting SApNP vaccines for both viruses (*58–60, 81*). It is therefore imperative to explore the cause(s) of spike metastability to facilitate rational vaccine design for SARS-CoV-2.

**Fig. 2.**
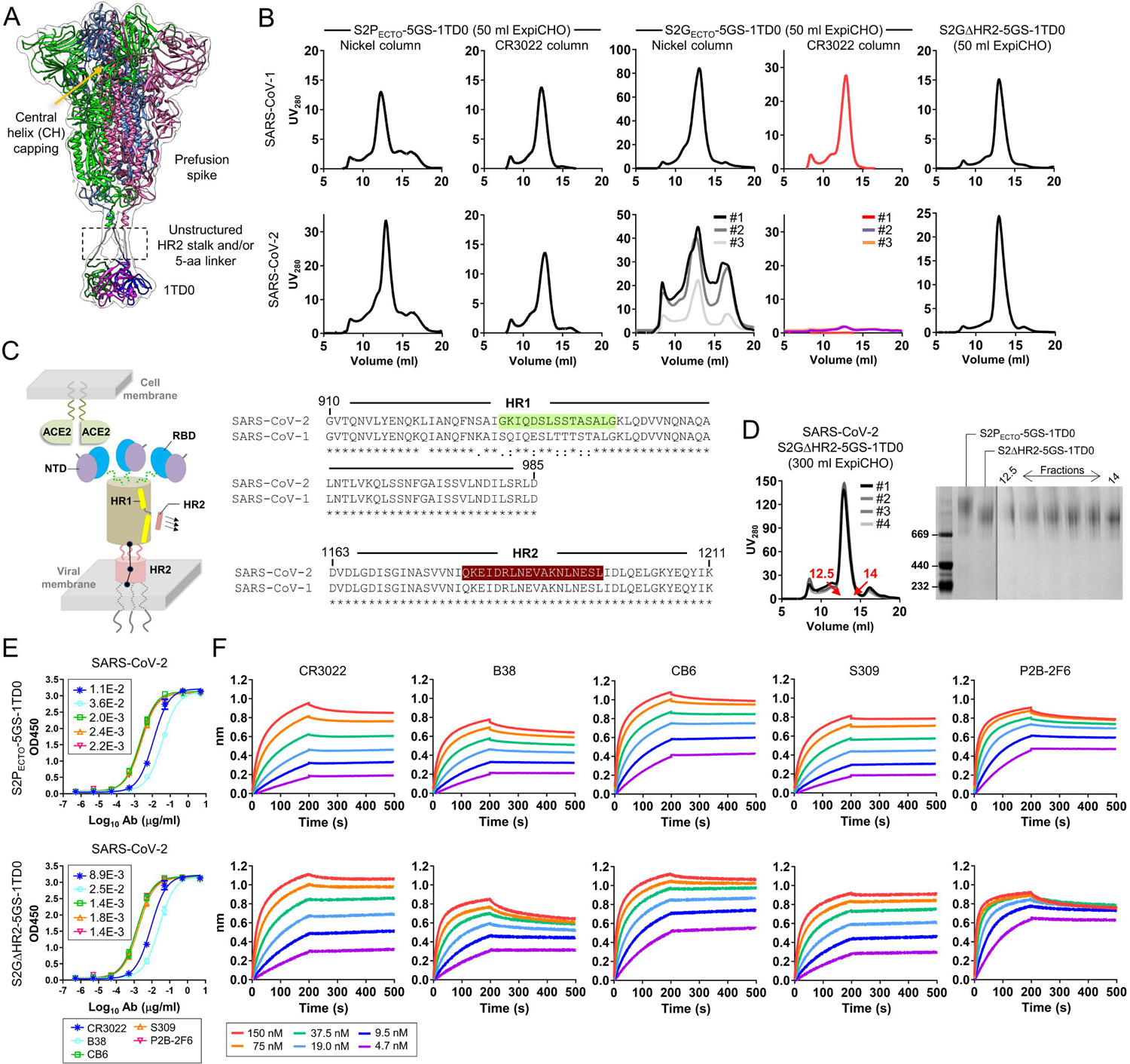
Rational design of SARS-CoV-2 spike antigens. **(A)** Structural model of prefusion S spike linked to the C-terminal trimerization domain (1TD0) with a 5GS linker in transparent molecular surface. The approximate position for the unstructured HR2 stalk, or in this case a 5-aa G_4_S linker, is highlighted with a dashed line box. **(B)** SEC profiles of SARS-CoV-1 (top) and SARS-CoV-2 (bottom) spikes. Left to right panels: S2P_ECTO_-5GS-1TD0 purified on Nickel and CR3022 columns (1 and 2), S2G_ECTO_-5GS-1TD0 on Nickel and CR3022 columns (3 and 4), and S2GΔHR2-5GS-1TD0 on a CR3022 column (5). All spike constructs tested in (B) contain a C-terminal His_6_ tag to facilitate Nickel column purification. Results from three separate production runs are shown for SARS-CoV-2 S2G_ECTO_-5GS-1TD0. **(C)** Schematic representation of a full-length SARS-CoV-2 spike on the virus membrane surface in the presence of host ACE2 and the HR2 region from a neighboring spike (left), and sequence alignment of SARS-CoV-1/2 HR1 and HR2 regions (right, top and bottom). The HR1 and HR2 segments that form a six-helix bundle in the post-fusion state are highlighted in green and brown shade, respectively. **(D)** Left: SEC profiles of S2GΔHR2-5GS-1TD0, showing results from four separate production runs. Right: BN-PAGE analysis of S2P_ECTO_-5GS-1TD0 and S2GΔHR2-5GS-1TD0. SEC fractions (12.5-14.0) are shown for S2GΔHR2-5GS-1TD0 on the gel. SEC profiles in (B) and (D) were obtained from a Superose 6 10/300 GL column. **(E)** ELISA binding of two SARS-CoV-2 spikes (S2P_ECTO_-5GS-1TD0 and S2GΔHR2-5GS-1TD0) to five mAbs. EC_50_ (μg/ml) values are labeled. **(F)** Octet binding of two SARS-CoV-2 spikes (S2P_ECTO_-5GS-1TD0 and S2GΔHR2-5GS-1TD0) to five mAbs. Sensorgrams were obtained from an Octet RED96 instrument at six antigen concentrations from 150 to 4.7 nM by twofold dilutions. The spike constructs tested in (D)-(F) do not contain a C-terminal His_6_ tag.

We first created uncleaved spike ectodomain (S_ECTO_) constructs for SARS-CoV-1/2, both containing the 2P mutation (K968P/V969P and K986P/V987P, respectively), a 5-aa G_4_S linker, a trimerization motif (PDB: 1TD0), and a C-terminal His_6_ tag. The two constructs were transiently expressed in 50-ml ExpiCHO cells followed by purification on either a Nickel column or a CR3022 column. The S2P_ECTO_-5GS-1TD0-His_6_ protein was characterized by SEC on a Superose 6 10/300 GL column (**Fig. 2B**, panels 1 and 2). After the Nickel column, both S2P_ECTO_ constructs showed a trimer peak (∼12 ml) with the left and right shoulders indicative of aggregate and dimer/monomer species, respectively. CR3022 purification resulted in a consistent trimer peak, as well as reduction in dimer/monomer species. We then compared a pair of S_ECTO_ constructs for SARS-CoV-1/2, both containing a double glycine (2G) mutation, K968G/V969G and K986G/V987G, respectively. The 2G mutation had little effect on the SARS-CoV-1 spike but produced abnormal SEC profiles and showed no yield for the SARS-CoV-2 spike after purification by Nickel and CR3022 columns, respectively (**Fig. 2B**, panels 3 and 4). Lastly, we tested a pair of S2G constructs without the HR2 stalk (E1150-Q1208), termed S2GΔHR2. Deletion of the HR2 stalk restored the SARS-CoV-2 trimer peak and reduced aggregates for both SARS-CoV-1/2, as shown by the SEC profiles upon CR3022 purification (**Fig. 2B**, panel 5). Since the triple-hinged HR2 stalk can generate diverse spike orientations on native virions (*54–56*), and the fusion core is formed by HR1 and HR2, we hypothesized that HR2 may be a key determinant of SARS-CoV-2 spike metastability (**Fig. 2C**, left). It is plausible that the interactions between HR1 and HR2 of two neighboring spikes may facilitate the pre-to-post-fusion transition in addition to ACE2 binding and S1 dissociation. Given the extensive sequence difference in HR1 (9 amino acids in total) compared to SARS-CoV-1 (**Fig. 2C**, right), we sought to examine the contribution of HR1 to SARS-CoV-2 spike metastability with two HR1-swapped (HR1_S_) spike constructs. Interestingly, while HR1 swapping proved ineffective, deletion of the HR2 stalk once again restored the trimer peak, suggesting a more important role for HR2 (**fig. S2**, **A** to **C**). Therefore, S2GΔHR2 appeared to be a general spike design for SARS-CoV-1/2 and perhaps other CoVs. Four separate production runs of SARS-CoV-2 S2GΔHR2-5GS-1TD0 in 300-ml ExpiCHO cells resulted in nearly identical SEC profiles with trimer yields of 0.8-1.0 mg (**Fig. 2D**, left). Consistently, blue native polyacrylamide gel electrophoresis (BN-PAGE) showed high trimer purity across SEC fractions (**Fig. 2D**, right). Antigenicity was assessed for CR3022/SEC-purified SARS-CoV-2 S2P_ECTO_ and S2GΔHR2 spike proteins. In ELISA, the S2GΔHR2 spike showed slightly higher affinity for the five representative mAbs than did the S2P_ECTO_ spike (**Fig. 2E**). When tested against three newly identified human NAbs, C105 (*49*) and CC12.1/CC12.3 (*41*), the two spikes yielded similar EC_50_ values (**fig. S2D**). In BLI, the S2GΔHR2 spike showed higher binding signals than the S2P_ECTO_ spike at the highest concentration, while exhibiting similar binding kinetics (**Fig. 2F**). The use of NAb P2B-2F6 (*36*) for spike purification resulted in higher trimer yield with similar purity to the CR3022 column across SEC fractions (**fig. S2E**). Thermostability was assessed by differential scanning calorimetry (DSC) for CR3022/SEC-purified SARS-CoV-2 S2P_ECTO_ and S2GΔHR2 spikes (**fig. S2F**). Upon the 2P-to-2G substitution and HR2 deletion, a small increase in thermal denaturation midpoint (T_m_) was observed (46.8 vs. 47.6 °C), with notably higher onset temperature (T_on_), 35.2 vs. 40.0 °C, and narrower half width of the peak (ΔT_1/2_), 4.7 vs. 3.9 °C. Altogether, we demonstrated that the HR2 stalk is a major source of spike metastability and S2GΔHR2 presents an alternative spike design to S2P, although more stabilizing mutations may be required to achieve greater thermostability.

### Rational design of single-component self-assembling spike nanoparticles

Although it was possible to conjugate trimeric SARS-CoV-2 spikes to an SApNP using the SPY system (*82*), the random, irreversible linking will likely result in irregular display with unoccupied but spatially occluded anchoring sites on the surface. The SPY system is perhaps more suitable for small individual antigens (e.g. the RBD). Using the gene fusion approach, we previously designed single-component SApNPs displaying stabilized HIV-1 Env trimers (*58, 59*) and optimized HCV E2 cores (*57*). Recently, we reengineered E2p and I3-01v9 60-mers to incorporate an inner layer of LDs to stabilize the non-covalently formed NP shell, in addition to a cluster of T-cell epitopes (*60*). This multilayered design may be essential to the stability of resulting SApNP vaccines when they are used to display large, complex viral antigens such as the SARS-CoV-2 spike.

Native SARS-CoV-2 virions present both pre- and post-fusion spikes (*52, 54, 55*) (**Fig. 3A**, top), while our vaccine strategy aims to develop single-component SApNPs that each present 8 or 20 stable prefusion S2GΔHR2 spikes to the immune system (**Fig. 3A**, bottom). As demonstrated in our previous studies (*58, 59*), different linker lengths may be needed to connect a trimer to the SApNP surface, as the spacing between the N termini of NP subunits around each three-fold axis varies: FR (20 Å), E2p (9 Å), and I3-01v9 (50.5 Å). Based on this consideration, we displayed the S2GΔHR2 spike on FR with a 5-aa G_4_S linker, on E2p with a 5-aa G_4_S linker, and on I3-01v9 with a 10-aa (G_4_S)_2_ linker, resulting in SApNPs with diameters of 47.9, 55.9, and 59.3 nm, respectively (**Fig. 3B**). The multilayered E2p-L4P and I3-01v9-L7P (*60*), which were validated for presenting RBDs (**Fig. 1**, **K** to **L**; **fig. S1D**), were used here as NP carriers of the S2GΔHR2 spike. Together, three S2GΔHR2 SApNP constructs were transiently expressed in 400-ml ExpiCHO cells, followed by CR3022 purification and SEC on a Superose 6 10/300 GL column (**Fig. 3C**). Three production runs for each of the three constructs generated highly consistent SEC profiles, despite the variation of low molecular weight (m.w.) impurities observed for the FR and E2p-L4P. After CR3022 and SEC purification, we obtained 0.3-0.4, 0.5-1.0, and 0.8-1.2 mg protein for S2GΔHR2-5GS-FR, S2GΔHR2-5GS-E2p-L4P, and S2GΔHR2-10GS-I3-01v9-L7P, respectively. Overall, the I3-01v9-derived S2GΔHR2 SApNP appeared to perform best in terms of particle yield, purity, and stability. The structural integrity was analyzed by nsEM, which showed well-formed particles of 45-65 nm with recognizable protrusions on the surface (**Fig. 3D**). The varying shape of these profusions may correspond to S2GΔHR2 spikes with “open” RBDs, in contrast to an array of “closed” HIV-1 and EBOV trimers on the SApNP surface (*58–60*). Such open conformation may facilitate induction of a strong RBD-specific NAb response *in vivo*. Antigenicity of S2GΔHR2 SApNPs was assessed using the same panel of mAbs. In ELISA, three SApNPs showed slightly improved binding to some, but not all, mAbs compared to the individual spike (**Fig. 3E**). In BLI, we observed a clear correlation between peak mAb binding signal and antigen valency, with E2p/I3-01v9 > FR > spike (**Fig. 3F**). Multivalent display on the two 60-mers substantially increased mAb binding compared to the FR 24-mer. In previous studies, we observed a similar correlation for HIV-1 Env trimer and HCV E2 core versus their SApNPs (*57, 58*). In summary, these VLP-size SApNPs with 8 or 20 spikes on the surface provide promising vaccine candidates for *in vivo* evaluation.

**Fig. 3.**
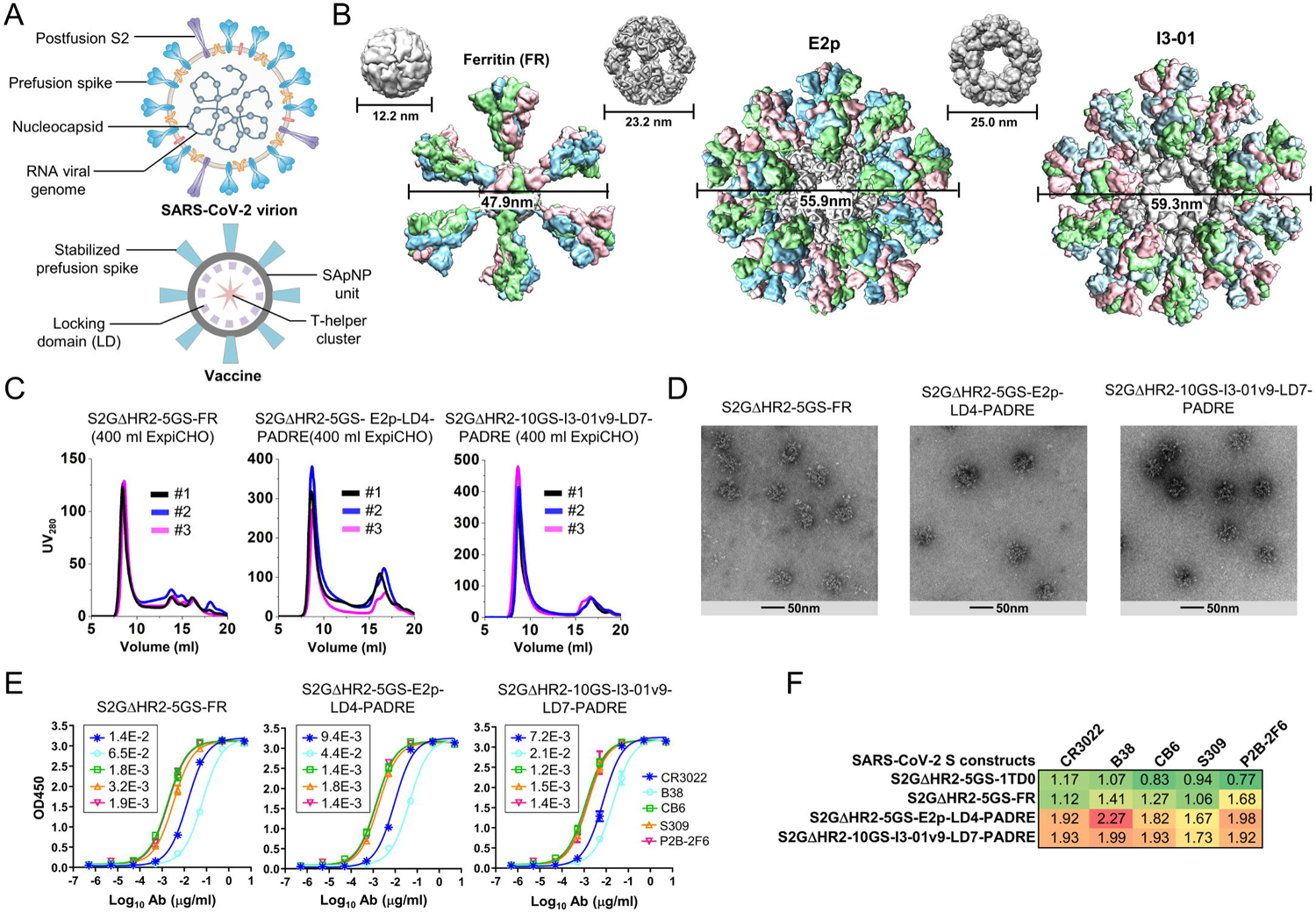
Rational design of SARS-CoV-2 spike-presenting nanoparticle vaccines. **(A)** Schematic representation of SARS-CoV-2 virion (top) and spike-presenting SApNP vaccine (bottom). For the SARS-CoV-2 virion, pre/post-fusion S, nucleocapsid, and RNA viral genome are shown, while for the vaccine, stabilized spike and multilayered SApNP composition are depicted. **(B)** Colored surface models of SApNP carriers (top) and spike-presenting SApNP vaccines (bottom). The three SApNP carriers used here are ferritin (FR) 24-mer and multilayered E2p and I3-01v9 60-mers (the LD layer and PADRE cluster are not shown). Particle size is indicated by diameter (in nanometers). **(C)** SEC profiles of three SApNPs presenting the SARS-CoV-2 S2GΔHR2 spike, from left to right, S2GΔHR2-5GS-FR, S2GΔHR2-5GS-E2p-LD4-PADRE (or E2p-L4P), and S2GΔHR2-10GS-I3-01v9-LD7-PADRE (or I3-01v9-L7P). SEC profiles were obtained from a Superose 6 10/300 GL column, each showing results from three separate production runs. **(D)** EM images of three SARS-CoV-2 S2GΔHR2 SApNPs. **(E)** ELISA binding of three SARS-CoV-2 S2GΔHR2 SApNPs to five mAbs. EC_50_ (μg/ml) values are labeled. **(F)** Antigenic profiles of SARS-CoV-2 S2GΔHR2 spike and three SApNPs against five mAbs. Sensorgrams were obtained from an Octet RED96 using six antigen concentrations (150-4.6 nM for spike, 9-0.27 nM for FR SApNP, and 3.5-0.1 nM for E2p and I3-01v9 SApNPs, respectively, all by twofold dilutions) and AHQ biosensors, as shown in fig. S3B. The peak binding signals (nm) at the highest concentration are listed. Color coding indicates the signal strength measured by Octet (green to red: low to high).

### SARS-CoV-1/2 vaccine-induced binding antibody response

Selected SARS-CoV-1/2 RBD- and spike-based vaccine constructs were assessed in BALB/c mice (**Fig. 4A**). We adopted a similar immunization protocol to be consistent with our previous studies on HIV-1, HCV, and EBOV SApNPs (*57, 58, 60*). Briefly, groups of five mice were immunized four times at three-week intervals. All antigens (50 μg per dose) were formulated with AddaVax, an oil-in-water emulsion adjuvant, except for I3-01v9, which was mixed with aluminum phosphate (AP) (*83*). We first analyzed the binding antibody (bAb) responses, as measured by EC_50_ titers, in the two SARS-CoV-2 RBD vaccine groups (**Fig. 4B** and **fig. S4**). The SPY-linked RBD SApNP (RBD-5GS-SPY-5GS-FR) elicited significantly higher bAb titers than the scaffolded RBD trimer (RBD-5GS-1TD0) at w2 and w5, irrespective of the coating antigen, and yielded a significant *P* value at w8 when the RBD was coated. Compared to the S2P_ECTO_ spike (S2P_ECTO_-5GS-1TD0), the RBD SApNP elicited significantly higher bAb titers against the RBD at w2, w5, and w8 (**Fig. 4B**, right), demonstrating a strong epitope-focusing effect. Mouse sera bound the SARS-CoV-1 spike with lower EC_50_ titers than the SARS-CoV-2 spike but with similar patterns (**fig. S4A**). We then analyzed the bAb responses induced by SARS-CoV-2 spikes S2P_ECTO_-5GS-1TD0 and S2GΔHR2-5GS-1TD0, as well as three SApNPs each displaying 8 or 20 S2GΔHR2 spikes (**Fig. 4C** and **fig. S5**). The S2GΔHR2 spike showed two-three-fold higher average EC_50_ titers than the S2P_ECTO_ spike irrespective of the coating antigen (of note, to facilitate a fair comparison, mouse sera from these two groups were tested against their respective spike antigens). Three SApNPs exhibited different temporal patterns depending on the coating antigen. In terms of spike-specific response, the I3-01v9 group registered a steady increase in average EC_50_ titer over time, showing the highest bAb titers at w2 and w8 and significantly outperforming the S2P_ECTO_ group at all time points. The I3-01v9 group also yielded higher EC_50_ titers than the S2GΔHR2 group throughout, although not with significant *P* values. The FR SApNP group exhibited a similar temporal pattern with lower EC_50_ titers, which were still significantly higher than the S2P_ECTO_ group. Among the three SApNPs, E2p exhibited the lowest average EC_50_ titer at w2 and reached the highest at w5, which then decreased slightly at w8. In terms of RBD-specific response, the five groups showed a clear ranking based on their average EC_50_ titers, which remained consistent across time points. At w2, I3-01v9 showed an average EC_50_ titer of 175, whereas all other spike-based vaccines induced little RBD-specific bAb response. At w5 and w8, S2GΔHR2 elicited higher bAb titers (on average by two-fold) than S2P_ECTO_, while all three SApNPs outperformed the individual S2GΔHR2 spike with a ranking of average EC_50_ titers correlated with their size (FR < E2p < I3-01v9). Sera reacted with the SARS-CoV-1 spike similarly, albeit at a lower level (**fig. S5A**). Lastly, we compared the bAb responses induced by three SARS-CoV-1 vaccines, S2P_ECTO_ spike (S2P_ECTO_-5GS-1TD0), scaffolded RBD trimer (RBD-5GS-1TD0), and SPY-linked RBD SApNP (RBD-5GS-SPY-5GS-FR) (**Fig. 4D** and **fig. S6**). Based on average EC_50_ titers, the SARS-CoV-1 S2P_ECTO_ spike appeared to have elicited a more robust bAb response than the SARS-CoV-2 S2GΔHR2 spike, whereas the SARS-CoV-1 RBD SApNP was less advantageous than its SARS-COV-2 counterpart. Serum reactivity with the SARS-CoV-2 S2P_ECTO_ spike was observed for all three SARS-CoV-1 vaccine groups (**fig. S6A**). In summary, RBD SApNPs can elicit higher titers of RBD-specific bAbs than the scaffolded RBD trimer and S2P_ECTO_ spike, albeit at different levels for the two SARS-CoVs. The S2GΔHR2 spike is more immunogenic than the S2P_ECTO_ spike, showing on average two-fold higher bAb titers. The S2GΔHR2-presenting E2p and I3-01v9 SApNPs are the best performers among all the spike-based vaccines, consistent with our previous findings for these two large SApNPs (*57, 58, 60*).

**Fig. 4.**
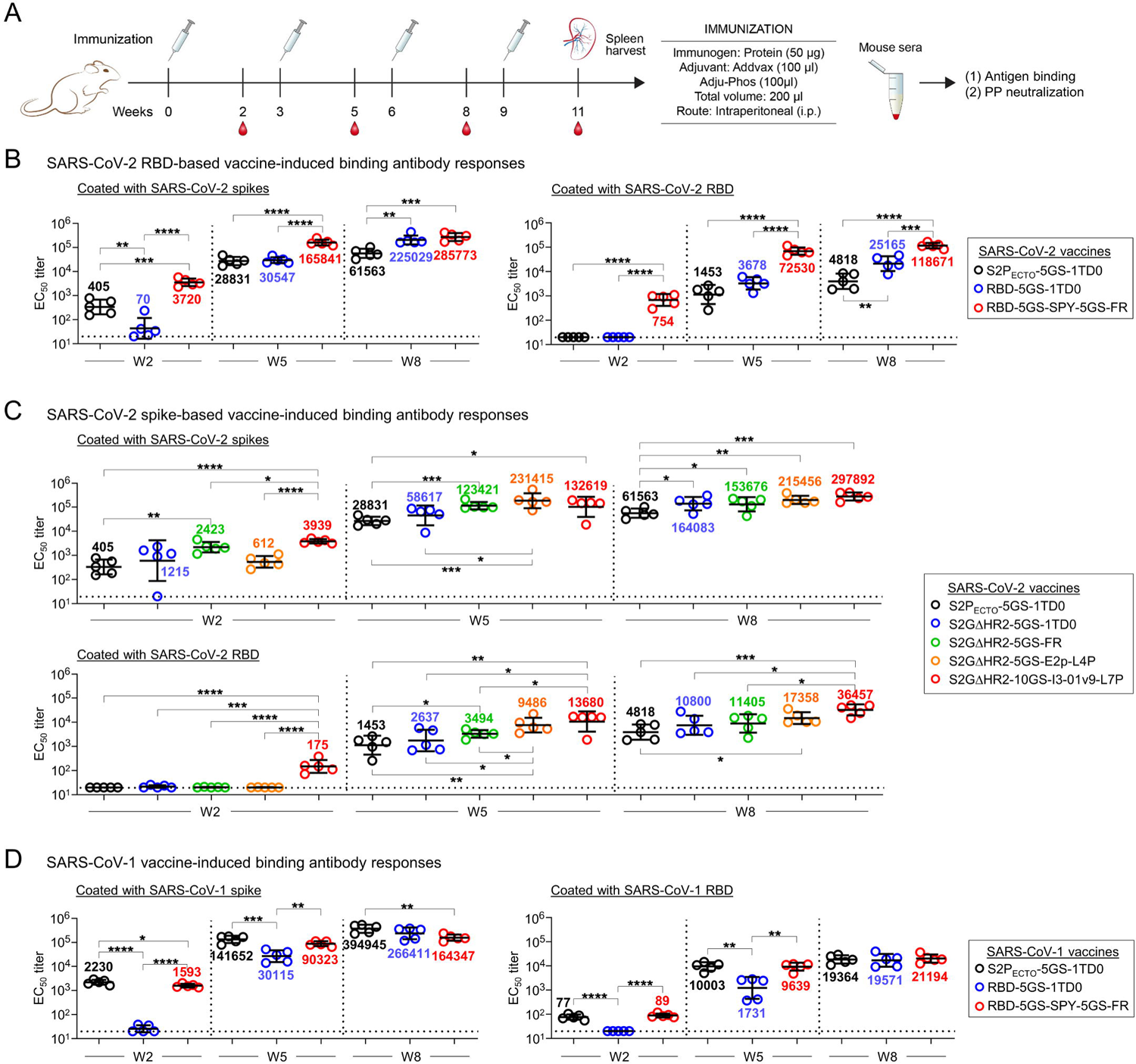
SARS-CoV-1/2 vaccine-induced binding antibody (bAb) responses in BALB/c mice. **(A)** Schematic representation of the mouse immunization protocol. **(B)** SARS-CoV-2 RBD and RBD-SApNP vaccine-induced bAb responses. Coating antigens: SARS-CoV-2 S2GΔHR2-5GS-foldon (left) and RBD (right). **(C)** SARS-CoV-2 S2GΔHR2 spike and S2GΔHR2-SApNP vaccine-induced bAb responses. Coating antigens: SARS-CoV-2 S2GΔHR2-5GS-foldon (top) and RBD (bottom). In (B) and (C), the SARS-CoV-2 S2P_ECTO_-5GS-1TD0 group is included for comparison, with S2P_ECTO_-5GS-foldon used as the coating antigen. **(D)** SARS-CoV-1 RBD and RBD-SApNP vaccine-induced bAb responses. Coating antigens: SARS-CoV-1 S2P_ECTO_-5GS-foldon (left) and RBD (right). In (D), the SARS-CoV-1 S2P_ECTO_-5GS-1TD0 group is included for comparison, with S2P_ECTO_-5GS-foldon used as the coating antigen. In (B) to (D), EC_50_ titers derived from the ELISA binding of mouse plasma to coating antigens are plotted, with average EC_50_ values labeled on the plots. The *P* values were determined by an unpaired *t* test in GraphPad Prism 8.4.3 with (*) indicating the level of statistical significance. Detailed ELISA data are shown in figs. S4-S6.

### SARS-CoV-1/2 vaccine-induced NAb response

One major goal in COVID-19 vaccine development is to generate a potent NAb response that can protect against SARS-CoV-2 infection. Pseudoparticle (SARS-CoV-1/2-pp) neutralization assays (*84*) were used to evaluate serum NAb responses elicited by different vaccines. We first analyzed the NAb responses, as measured by 50% inhibitory dilution (ID_50_) titers, in the two SARS-CoV-2 RBD vaccine groups (**Fig. 5A** and **fig. S7**). The SPY-linked RBD SApNP elicited a NAb response against autologous SARS-CoV-2 as early as w2, albeit with low titers, and retained its advantage at w5 and w8, suggesting that such RBD SApNP vaccines can elicit a rapid NAb response. The scaffolded RBD trimer group showed higher average ID_50_ titers than the S2P_ECTO_ spike group at both w5 and w8. A somewhat different pattern was observed in the SARS-CoV-1-pp assay. At w2, no vaccine group showed detectable heterologous NAb response. At w5 and w8, the S2P_ECTO_ spike elicited a more potent anti-SARS-CoV-1 NAb response than the scaffolded RBD trimer, suggesting that non-RBD epitopes on the spike may contribute to cross-neutralization. We then analyzed the NAb responses induced by five spike-based vaccines (**Fig. 5B** and **fig. S8**). In terms of autologous neutralization, no spike-based vaccine elicited any NAb response that blocks SARS-CoV-2-pps after one dose, but a consistent pattern of serum neutralization was observed at w5 and w8 (**Fig. 5B**, upper panel). Specifically, the S2P_ECTO_ spike showed the lowest average ID_50_ titers, 879 and 2481 at w5 and w8, respectively, whereas the S2GΔHR2 spike induced a stronger NAb response with 2.8-6.7-fold higher average ID_50_ titers, which did not reach *P* ≤ 0.05 due to within-group variation. Nonetheless, this result confirmed the beneficial effect of the 2G mutation and the HR2 stalk deletion on NAb elicitation. Among the three SApNPs, E2p was the best performer at w5, showing an average ID_50_ titer of 8435 that is 9.6-fold higher than S2P_ECTO_ and 1.4-fold higher than S2GΔHR2, while I3-01v9 showed the most potent NAb response at w8 with an average ID_50_ titer of 17351 that is about 7-fold and 2.5-fold higher than S2P_ECTO_ and S2GΔHR2, respectively. A similar temporal pattern was observed in the heterologous SARS-CoV-1-pp assay (**Fig. 5B**, lower panel). It is worth noting that the multilayered I3-01v9 SApNP elicited a SARS-CoV-1 NAb response with an average ID_50_ titer of 351 at w2, whereas all other groups showed no detectable serum neutralization. These results suggest that a well-designed SARS-CoV-2 S2GΔHR2 SApNP vaccine may provide protection against both SARS-CoVs. Lastly, we analyzed the NAb responses induced by three SARS-CoV-1 vaccines (**Fig. 5C** and **fig. S9**). In the autologous SARS-CoV-1-pp assay, the S2P_ECTO_ spike and the RBD SApNP induced significantly higher NAb titers than the scaffolded RBD trimer at w2 and w5, and all three vaccine groups showed similar ID_50_ titers at w8. However, heterologous SARS-CoV-2 neutralization was below or at the baseline level for three SARS-CoV-1 vaccines at w2, w5, and w8. In this study, the pseudovirus neutralization assay was validated using a panel of known SARS-CoV-1/2 NAbs (**fig. S9C**). As a negative control, the w8 mouse sera were tested against pseudoparticles bearing the murine leukemia virus (MLV) Env, MLV-pps, showing no detectable reactivity (**fig. S9D**). In summary, these results demonstrate an advantage in NAb elicitation by the SARS-CoV-2 S2GΔHR2 spike and its SApNPs compared to the widely used S2P_ECTO_ spike. Although SARS-CoV-2 RBD- and S2GΔHR2-presenting SApNPs are both effective at eliciting an autologous NAb response, the latter can induce high NAb titers to SARS-CoV-1 and may provide broader protection against SARS-associated CoVs.

**Fig. 5.**
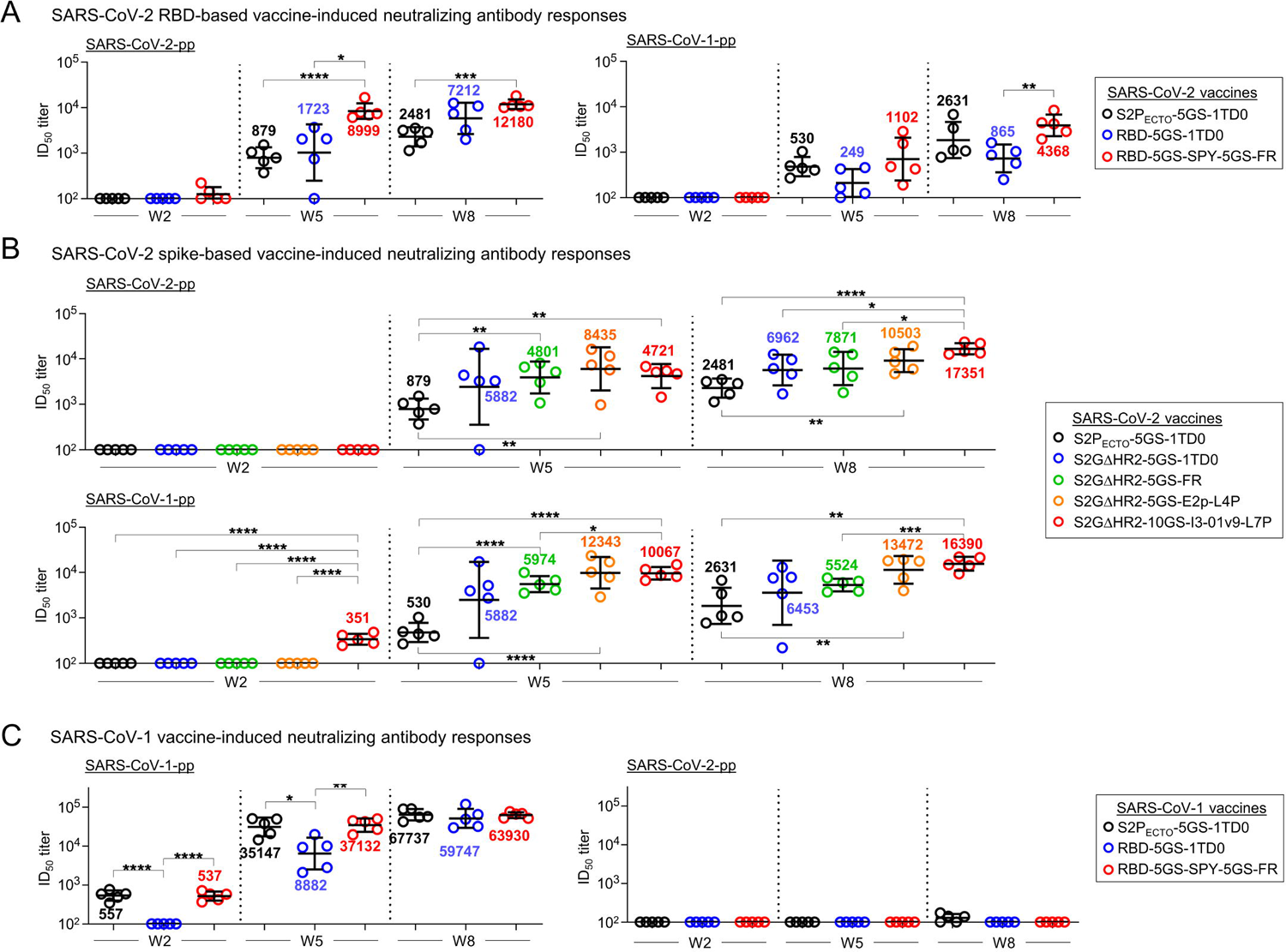
SARS-CoV-1/2 vaccine-induced neutralizing antibody (NAb) responses in BALB/c mice. **(A)** SARS-CoV-2 RBD and RBD-SApNP vaccine-induced NAb responses to autologous SARS-CoV-2 (left) and heterologous SARS-CoV-1 (right). **(B)** SARS-CoV-2 S2GΔHR2 spike and S2GΔHR2-SApNP vaccine-induced NAb responses to SARS-CoV-2 (top) and SARS-CoV-1 (bottom). In (A) and (B), the SARS-CoV-2 S2P_ECTO_-5GS-1TD0 group is included for comparison. **(C)** SARS-CoV-1 RBD and RBD-SApNP vaccine-induced NAb responses to autologous SARS-CoV-1 (left) and heterologous SARS-CoV-2 (right). In (C), the SARS-CoV-1 S2P_ECTO_-5GS-1TD0 group is included for comparison. In (A) to (C), ID_50_ titers derived from the SARS-CoV-1/2-pp neutralization assays are plotted, with average ID_50_ values labeled on the plots. The *P* values were determined by an unpaired *t* test in GraphPad Prism 8.4.3 with (*) indicating the level of statistical significance. Detailed SARS-CoV-1/2-pp neutralization data are shown in figs. S7-S9.

### SARS-CoV-2 vaccine-induced T-cell response

While humoral immunity is required to block host-virus interaction and prevent viral infection, cellular immunity is essential for eliminating infected host cells to control viral infection (*85–88*). Emerging evidence indicates that an early T-cell response (*89, 90*), as well as T-cell memory (*91*), is critical for protection against SARS-CoV-2. However, COVID-19 vaccines must induce a CD4^+^ T helper 1 (Th1), but not Th2-type, T-cell response, as the latter has been implicated in vaccine-associated enhancement of respiratory disease (VAERD) (*10*). In addition, T follicular helper cells (Tfh) play an important role in the maturation and production of NAbs. Therefore, understanding T-cell responses is crucial for development of an effective and safe COVID-19 vaccine.

Interferon (IFN)-γ-producing Th1 cells are important for generating an optimal antibody response and for induction of cellular immunity (*85–87*). We first examined various SARS-CoV-2 vaccines on induction of CD4^+^ Th1 responses specific to the vaccine antigen at w11, two weeks after the fourth immunization, when memory T cells had already developed in spleen (*88*). Mouse splenocytes from the S2P_ECTO_ group and two S2GΔHR2 SApNP groups (E2p and I3-01v9) were analyzed by flow cytometry using naïve samples as a negative control. The I3-01v9 group induced about 1.5- and 2.3-fold higher frequency of IFN-γ-producing CD4^+^ Th1 cells than the S2P_ECTO_ and E2p groups, respectively (**Fig. 6A**). Notably, after re-stimulation with the respective antigens for as few as 4 hours, both SApNP groups produced ∼two-fold higher frequency of CD107a-producing cytolytic CD4^+^ T cells than the S2P_ECTO_ and naïve groups (**Fig. 6B**). IFN-γ/IL-4 (interleukin-4) double-positive cells are memory CD4^+^ T cells that have acquired the ability to produce IL-4 while still retaining the ability to produce IFN-γ under Th1 conditions (*92*). It appeared that I3-01v9 induced three- and five-fold more IFN-g/IL-4 double-positive memory CD4^+^ T cells than S2P_ECTO_ and E2p (**Fig. 6A**). These results suggest that I3-01v9 can induce both CD4^+^ Th1 cells and IFN-γ/IL-4 double-positive memory CD4^+^ T cells. In addition, I3-01v9 induced more IFN-γ/GM-CSF (granulocyte-macrophage colony-stimulating factor) double-positive CD8^+^ effector T cells than S2P_ECTO_ and E2p (**Fig. 6C**), suggesting that protective CD8^+^ T cells were also generated in mice immunized with I3-01v9. Of note, CD8^+^ T cells derived from mice immunized with I3-01v9, rather than those with S2P_ECTO_ and E2p, acquired the ability to rapidly produce IFN-γ upon antigen re-stimulation (**Fig. 6D**), suggesting generation of I3-01v9-responsive effector/memory T cells. Our results indicate that the S2GΔHR2 I3-01v9 SApNP can induce robust T-cell responses consisting of CD4^+^ Th1 cells, IFN-γ/IL-4 double-positive memory CD4^+^ T cells, and effector CD8^+^ T cells, thus providing protective cellular immunity in addition to a potent NAb response. Since T cell immunity against the SApNP backbone cannot be ruled out, a more detailed T-cell analysis using soluble spikes, spike peptides, and naked SApNPs for re-stimulation may be warranted.

**Fig. 6.**
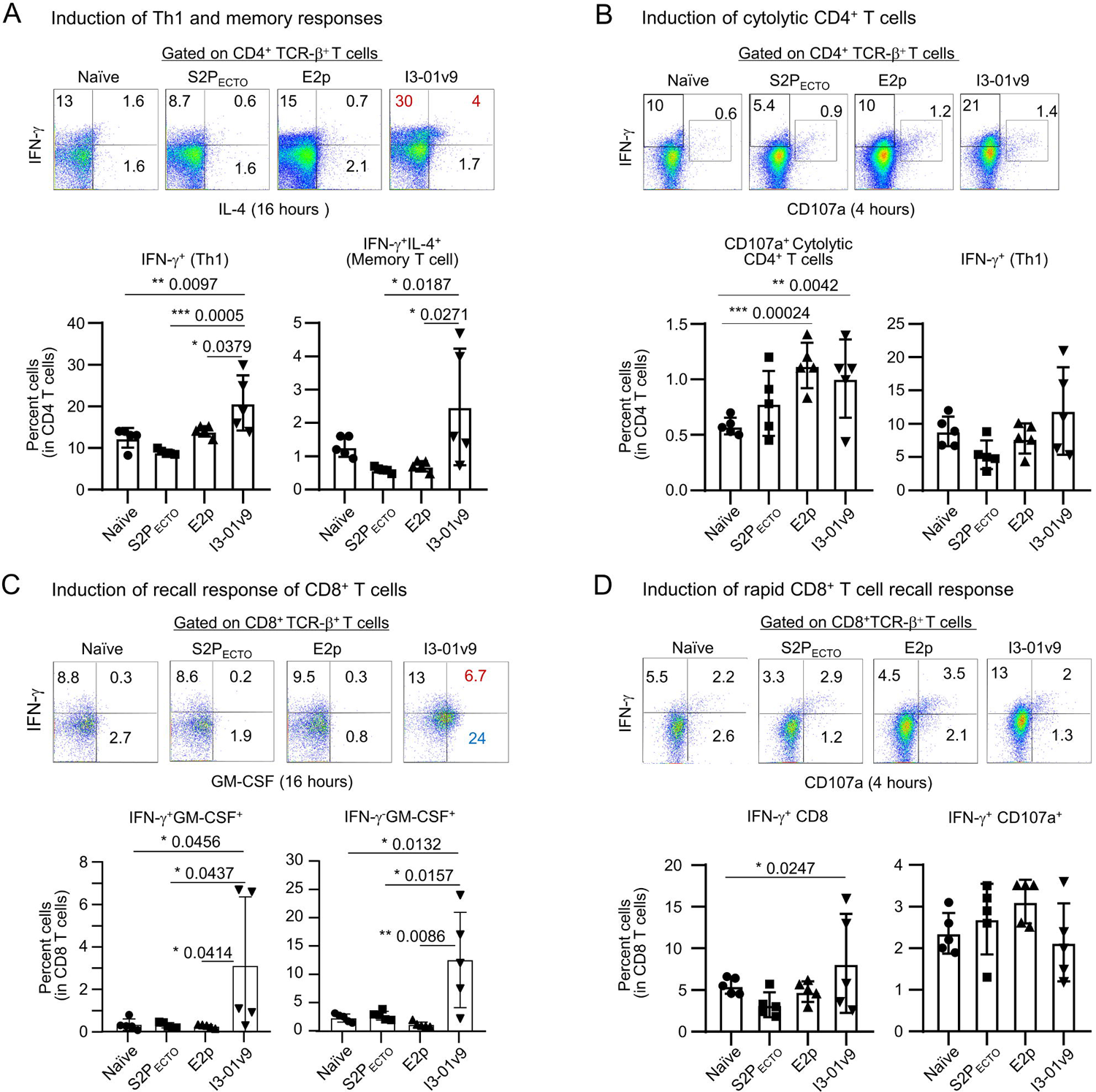
SARS-CoV-2 vaccine-induced T-cell responses in BALB/c mice. Splenocytes from mice (n=5 in each group) immunized with S2P_ECTO_ spike, S2GΔHR2 E2p SApNP, and S2GΔHR2 I3-01v9 SApNP, were isolated at w11 and cultured in the presence of IL-2 and dendritic cells (DC) pulsed with S2P_ECTO_ (1×10^-7^ μM), S2GΔHR2 E2p (1×10^-7^ μM), and S2GΔHR2 I3-01v9 (1×10^-7^ μM), respectively. Splenocytes from 5 naïve mice were used as control samples and cultured in the presence of DCs without antigen pulsing. Cells were assessed after 16 hours (A, C) and 4 hours (B, D) of culture. **(A)** and **(B)**: Vaccine-induced CD4^+^ T cell immunity. **(C)** and **(D)**: Vaccine-induced CD8^+^ T cell immunity. Plots show the frequencies of cell fraction. The *P* values were determined by one-way ANOVA analysis. *, *P* < 0.05; **, *P* < 0.01; ***, *P* < 0.001.

## DISCUSSION

COVID-19 is the first worldwide pandemic of this scale since the infamous Spanish influenza over a century ago (*93*), which caused ∼50 million deaths worldwide and remains a painful reminder of our vulnerability to a new virus without a protective vaccine. The rapid spread of SARS-CoV-2 therefore demanded rapid vaccine development (*10*). Operation Warp Speed (OWS) was launched in May 2020 with the initial goal to deliver 300 million doses of safe and effective vaccines by January 2021 (*94*), although that goal has not yet been realized. Nevertheless, in December 2020, while other vaccine candidates were still being evaluated in clinical trials, two OWS-supported mRNA vaccines were approved for EUA, marking an important turning point in the battle against the pandemic. However, vaccine development during a pandemic against a new virus poses unique challenges, one of which is how to balance public health needs versus scientific rigor (*95–97*). The global vaccine campaign also provides a unique opportunity to compare different vaccine design strategies and platforms – especially new ones – against a common target. While mRNA and viral vector vaccines remain the front runners, protein-based vaccines are highlighted in a recent review with the prediction that they will eventually reach a larger fraction of the global population (*98*). As the NAb titers induced by the first-generation nucleic acid vaccines wane over time, effective protein vaccines will be needed to sustain long-term immunity against SARS-CoV-2.

Here, we approached SARS-CoV-2 vaccine with a rational design strategy and set out to develop protein nanoparticle vaccines that can be used alone or as a booster vaccine. To this end, a panel of vaccine candidates have been generated based on three SApNP platforms with different *in vitro* and *in vivo* attributes (**table S1**). Our comparative analysis of these vaccine constructs has offered some valuable insights. First, the choice of antigen is critical to the success of SARS-CoV-2 vaccine irrespective of the delivery platform. Most vaccine antigens including OWS’s vaccine candidates are based on S2P, which produces a spike structure that differs in detail from the full-length wildtype spike, e.g. in FPPR of S2 and in the relative dispositions of the S1 domains (*52*). These differences may complicate interpretation of vaccine outcome. S2P and other empirical spike designs (*53*) have attempted to constrain the spike conformation and increase trimer yield. However, as we previously demonstrated for HIV-1 Env and EBOV GP (*58, 60, 81*), it is important to identify and eliminate (if possible) the cause of spike metastability. During antigen screening, we found that deletion of the HR2 stalk with a 2P-to-2G substitution renders a more stable spike, which is consistent with recent reports on a highly flexible HR2 stalk in the native spikes on SARS-CoV-2 virions (*54–56*). Thus, S2GΔHR2 would seem to represent a major advance in spike design. Second, single-component SApNPs provide a powerful platform for vaccine development against diverse viral pathogens (*57, 58, 60*). Here, S2GΔHR2 was genetically fused, rather than chemically linked, to three SApNPs, including two multilayered 60-meric carriers with enhanced stability and an embedded T-help signal. Such protein vaccines should be more effective in eliciting a potent NAb response and less likely to induce adverse responses (*98, 99*). An epitope-focused vaccine strategy was also explored by designing scaffolded RBD trimers and SPY-linked RBD SApNPs. Third, to achieve high efficacy and ensure safety, vaccine-induced NAb and T-cell responses must be evaluated in animals prior to clinical trials. Indeed, in our mouse study, S2GΔHR2 appeared to be more effective than S2P in NAb elicitation, both alone and displayed on SApNPs. Of note, the multilayered S2GΔHR2 I3-01v9 SApNP elicited not only high NAb titers but also desired T-cell responses. In addition to CD4^+^ Th1 cells and memory CD4^+^ T cells, this SApNP also induced CD107a-producing cytolytic CD4^+^ T cells, which may directly kill infected host cells, and GM-CSF-producing CD8^+^ effector T cells, which may promote the generation of macrophages and functional dendritic cells (DCs) to facilitate the clearance of infected cells. It is also worth noting that the NAb response to the large SApNPs plateaued after three doses, suggesting that the number of injections and dosage can be reduced without diminishing vaccine-induced responses.

Lastly, expression of vaccine antigens in CHO cells followed by purification using an antibody column, such as CR3022, would allow rapid and industrial-scale vaccine production. In summary, our study provides promising COVID-19 vaccine candidates for evaluation in clinical trials.

## MATERIALS AND METHODS

### Design, expression, and purification of SARS-CoV-1/2 RBD and spike antigens

The spike (S) genes of the SARS-CoV-1 isolate Tor2 (GenBank accession #: NC_004718) and the SARS-CoV-2 isolate Wuhan-Hu-1 (GenBank accession #: MN908947) were used to design all the RBD and spike constructs following codon-optimization for expression in mammalian cells. The RBD sequence is defined as P317-D518 and P330-N532 for SARS-CoV-1 and 2, respectively. The S_ECTO_ sequence is defined as M1-Q1190 and M1-Q1208 for SARS-CoV-1 and 2, respectively. To remove the S1/S2 cleavage site, an R667G mutation and a ^682^GSAGSV^687^ modification were introduced in the SARS-CoV-1 and 2 spikes, respectively. The 2P (or 2G) mutation was made to K968/V969 and K986/V987 in the SARS-CoV-1 and 2 spikes, respectively. The SARS-CoV-2 C-terminal region (E1150-Q1208) containing the HR2 stalk was removed from S2G_ECTO_, resulting in an HR2-deleted spike construct, termed S2GΔHR2. The viral capsid protein SHP (PDB: 1TD0) was used as a trimerization motif in spike constructs for immunization, whereas the foldon domain from the bacteriophage T4 fibritin (PDB: 1RFO) was used in coating spike antigens for ELISA to mask the 1TD0-derived antibody response. All constructs were transiently expressed in ExpiCHO cells (Thermo Fisher). Briefly, ExpiCHO cells were thawed and incubated with ExpiCHO^TM^ Expression Medium (Thermo Fisher) in a shaker incubator at 37 °C, 135 rpm and 8% CO_2_. When the cells reached a density of 10×10^6^ ml^-1^, ExpiCHO^TM^ Expression Medium was added to reduce cell density to 6×10^6^ ml^-1^ for transfection. The ExpiFectamine^TM^ CHO/plasmid DNA complexes were prepared for 100-ml transfection in ExpiCHO cells following the manufacturer’s instructions. For a given construct, 100 μg of plasmid and 320 μl of ExpiFectamine CHO reagent were mixed in 7.7 ml of cold OptiPRO™ medium (Thermo Fisher). After the first feed on day one, ExpiCHO cells were cultured in a shaker incubator at 33 °C, 115 rpm and 8% CO_2_ following the Max Titer protocol with an additional feed on day five (Thermo Fisher). Culture supernatants were harvested 13 to 14 days after transfection, clarified by centrifugation at 4000 rpm for 25 min, and filtered using a 0.45 μm filter (Thermo Fisher). The CR3022 antibody column was used to extract SARS-CoV-1/2 antigens from the supernatants, which was followed by SEC on a Superdex 200 10/300 GL column (for scaffolded RBD trimers) or a Superose 6 10/300 GL column (for SPY-linked RBD SApNPs, spikes, and S2GΔHR2 SApNPs). A human NAb, P2B-2F6 (*36*), was also used to pack antibody columns for purification of the SARS-CoV-2 S2GΔHR2 spike, which was followed by SEC on a HiLoad 16/600 Superose 6 column. For comparison, His_6_-tagged S_ECTO_-5GS-1TD0 spike protein was extracted from the supernatants using an immobilized Ni Sepharose^TM^ Excel column (GE Healthcare) and eluted with 500 mM Imidazole prior to SEC. Protein concentration was determined using UV_280_ absorbance with theoretical extinction coefficients.

### Blue native polyacrylamide gel electrophoresis

SARS-CoV-2 spikes and spike-presenting SApNPs were analyzed by blue native polyacrylamide gel electrophoresis (*BN-PAGE*) and stained with Coomassie blue. The proteins were mixed with^TM^ sample buffer and G250 loading dye and added to a 4-12% Bis-Tris NativePAGE gel (Life Technologies). BN-PAGE gels were run for 2 to 2.5 hours at 150 V using the NativePAGE^TM^ running buffer (Life Technologies) according to the manufacturer’s instructions.

### Enzyme-linked immunosorbent assay

Each well of a Costar^TM^ 96-well assay plate (Corning) was first coated with 50 µl PBS containing 0.2 μg of the appropriate antigens. The plates were incubated overnight at 4 °C, and then washed five times with wash buffer containing PBS and 0.05% (v/v) Tween 20. Each well was then coated with 150 µl of a blocking buffer consisting of PBS and 40 mg ml^-1^ blotting-grade blocker (Bio-Rad). The plates were incubated with the blocking buffer for 1 hour at room temperature, and then washed five times with wash buffer. For antigen binding, antibodies (in the immunoglobulin G (IgG) form) were diluted in the blocking buffer to a maximum concentration of 5 μg ml^-1^ followed by a 10-fold dilution series. For each antibody dilution, a total of 50 μl volume was added to the appropriate wells. For mouse sample analysis, plasma was diluted by 20-fold in the blocking buffer and subjected to a 10-fold dilution series. For each sample dilution, a total of 50 μl volume was added to the wells. Each plate was incubated for 1 hour at room temperature, and then washed 5 times with PBS containing 0.05% Tween 20. For antibody binding, a 1:5000 dilution of goat anti-human IgG antibody (Jackson ImmunoResearch Laboratories, Inc), or for mouse sample analysis, a 1:3000 dilution of horseradish peroxidase (HRP)-labeled goat anti-mouse IgG antibody (Jackson ImmunoResearch Laboratories), was then made in the wash buffer (PBS containing 0.05% Tween 20), with 50 μl of this diluted secondary antibody added to each well. The plates were incubated with the secondary antibody for 1 hour at room temperature, and then washed 6 times with PBS containing 0.05% Tween 20. Finally, the wells were developed with 50 μl of TMB (Life Sciences) for 3-5 min before stopping the reaction with 50 μl of 2 N sulfuric acid. The resulting plate readouts were measured at a wavelength of 450 nm. Of note, the w2 serum binding did not reach the plateau (or saturation) to allow for accurate determination of EC_50_ titers. Nonetheless, the EC_50_ values at w2 were derived by setting the lower/upper constraints of OD_450_ at 0.0/3.2 to facilitate the comparison of different vaccine groups at the first time point.

### Bio-layer interferometry

The kinetics of SARS-CoV-1/2 vaccine antigens, RBD versus RBD-presenting SApNPs as well as spike versus spike-presenting SApNPs, binding to a panel of known antibodies (in the IgG form) was measured using an Octet RED96 instrument (FortéBio, Pall Life Sciences). All assays were performed with agitation set to 1000 rpm in FortéBio 1× kinetic buffer. The final volume for all the solutions was 200 μl per well. Assays were performed at 30 °C in solid black 96-well plates (Geiger Bio-One). For all antigens except for S2GΔHR2 SApNPs, 5 μg ml of antibody in 1× kinetic buffer was loaded onto the surface of anti-human Fc Capture Biosensors (AHC) for 300 s. For S2GΔHR2 SApNPs, anti-human Fc Quantitation Biosensors (AHQ) were used. A 60 s biosensor baseline step was applied prior to the analysis of the association of the antibody on the biosensor to the antigen in solution for 200 s. A two-fold concentration gradient of antigen, starting at 950 nM for scaffolded RBD trimers, 37 nM for SPY-linked RBD FR SApNP (RBD-5GS-SPY-5GS-FR), 150 nM for soluble spikes, and 9/3.5/3.5 nM for S2GΔHR2 presented on FR/E2p/I3-01v9 SApNPs, was used in a titration series of six. The dissociation of the interaction was followed for 300 s. Correction of baseline drift was performed by subtracting the mean value of shifts recorded for a sensor loaded with antibody but not incubated with antigen, and for a sensor without antibody but incubated with antigen. Octet data were processed by FortéBio’s data acquisition software v.8.1. Experimental data were fitted with the binding equations describing a 2:1 interaction to achieve optimal fitting. Of note, the S2GΔHR2 spike was also measured using AHQ biosensors to facilitate comparison of mAb binding with the S2GΔHR2-presenting SApNPs.

### Differential scanning calorimetry (DSC)

Thermal melting curves of SARS-CoV-2 S2_PECTO_-5GS-1TD0 and S2GΔHR2-5GS-1TD0 spike proteins following CR3022 and SEC purification were obtained from a MicroCal PEAQ-DSC Man instrument (Malvern). The purified spike trimer protein produced from ExpiCHO cells were buffer exchanged into 1×PBS and concentrated to 0.8 μM before analysis by the instrument. Melting was probed at a scan rate of 60 °C·h^−1^ from 20 °C to 100 °C. Data processing, including buffer correction, normalization, and baseline subtraction, was conducted using the MicroCal PEAQ-DSC software. Gaussian fitting was performed using the Origin 9.0 software.

### Electron microscopy (EM) assessment of nanoparticle constructs

The EM analysis of various RBD and S2GΔHR2-presenting SApNPs was performed by the Core Microscopy Facility at The Scripps Research Institute. All SApNP samples were prepared at the concentration of 0.01-0.05 mg/ml. Carbon-coated copper grids (400 mesh) were glow-discharged and 8 µL of each sample was adsorbed for 2 min. Excess sample was wicked away and grids were negatively stained with 2% uranyl formate for 2 min. Excess stain was wicked away and the grids were allowed to dry. Samples were analyzed at 80 kV with a Talos L120C transmission electron microscope (Thermo Fisher) and images were acquired with a CETA 16M CMOS camera.

### Animal immunization and sample collection

Similar immunization protocols have been reported in our previous vaccine studies (*57, 58, 60*). Briefly, the Institutional Animal Care and Use Committee (IACUC) guidelines were followed with animal subjects tested in the immunization study. Eight-week-old BALB/c mice were purchased from The Jackson Laboratory and housed in ventilated cages in environmentally controlled rooms at The Scripps Research Institute, in compliance with an approved IACUC protocol and AAALAC (Association for Assessment and Accreditation of Laboratory Animal Care) International guidelines. Mice were immunized at weeks 0, 3, 6, and 9 with 200 μl of antigen/adjuvant mix containing 50 μg of vaccine antigen and 100 μl of adjuvant, AddaVax or Adju-Phos (InvivoGen), via the intraperitoneal (i.p.) route. Blood was collected two weeks after each immunization. All bleeds were performed through the retro-orbital sinus using heparinized capillary tubes into EDTA-coated tubes. Samples were spun at 1200 RPM for 10 min to separate plasma (top layer) and the rest of the whole blood layer. Upon heat inactivation at 56 °C for 30 min, the plasma was spun at 2000 RPM for 10 min to remove precipitates. The rest of the whole blood layer was diluted with an equal volume of PBS and then overlaid on 4.5 ml of Ficoll in a 15 ml SepMate^TM^ tube (STEMCELL Technologies) and spun at 1200 RPM for 10 min at 20 °C to separate peripheral blood mononuclear cells (PBMCs). Cells were washed once in PBS and then resuspended in 1 ml of ACK Red Blood Cell lysis buffer (Lonza). After washing with PBS, PBMCs were resuspended in 2 ml of Bambanker Freezing Media (Lymphotec). Spleens were harvested at w11 and ground against a 70-μm cell strainer (BD Falcon) to release the splenocytes into a cell suspension. Splenocytes were centrifuged, washed in PBS, treated with 5 ml of ACK lysing buffer (Lonza), and frozen with 3ml of Bambanker freezing media. Plasma was used for ELISA and neutralization assays to determine binding and neutralizing antibody responses in mouse serum.

### SARS-CoV-1/2 pseudovirus neutralization assay

Pseudoparticle (SARS-CoV-1/2-pp) neutralization assays were utilized to assess the neutralizing activity of previously reported antibodies and vaccine-induced murine antibody response. SARS-CoV-1/2-pps were generated by co-transfection of HEK293T cells with the HIV-1 pNL4-3.lucR-E-plasmid (obtained from the NIH AIDS reagent program: https://www.aidsreagent.org/) and the expression plasmid encoding the S gene of SARS-CoV-1 isolate Tor2 (GenBank accession #: NC_004718) and the SARS-CoV-2 isolate Wuhan-Hu-1 (GenBank accession #: MN908947) at a 4:1 ratio by lipofectamine 3000 (Thermo Fisher Scientific). After 48 to 72 hours, SARS-CoV-1/2-pps were collected from the supernatant by centrifugation at 4000 rpm for 10 min, aliquoted, and stored at −80 °C before use. The mAbs at a starting concentration of 0.1-10 μg/ml, or mouse plasma at a starting dilution of 100-fold, were mixed with the supernatant containing SARS-CoV-1/2-pps and incubated for 1 hour at 37°C in white solid-bottom 96-well plate (Corning). A 3-fold dilution series was used in the assay. The HEK293T-hACE2 cell line (catalogue#: NR-52511) and the vector pcDNA3.1(-) containing the SARS-CoV-2 S gene (catalogue#: NR52420) were obtained from BEI RESOURCES (https://www.beiresources.org/) and used in pseudovirus neutralization assays (*84*). Briefly, HEK293T-hACE2 cells at 1×10^4^ were added to each well and the plate was incubated at 37°C for 48 hours. After incubation, overlying media was removed, and cells were lysed. The firefly luciferase signal from infected cells was determined using the Bright-Glo Luciferase Assay System (Promega) according to the manufacturer’s instructions. Data were retrieved from a BioTek microplate reader with Gen 5 software, the average background luminescence from a series of uninfected wells was subtracted from each well, and neutralization curves were generated using GraphPad Prism 8.4.3, in which values from wells were compared against a well containing SARS-CoV-1/2-pp only. The same HIV-1 vectors pseudotyped with the murine leukemia virus (MLV) Env gene, termed MLV-pps, were produced in HEK293T cells and included in the neutralization assays as a negative control. As the NAb titers plateaued after w8, the w11 results were not shown in Fig. 5 but included in figs. S7-S9 for comparison.

### Dendritic cell (DC) production

Mouse bone marrow (BM) was cultured in RPMI 1640 medium containing 10% fetal bovine serum and recombinant mouse Flt3L (50 ng/mL) and SCF (10 ng/ml) for 9 days as described (*100*). To induce DC activation, immature DCs were incubated with lipopolysaccharide (LPS, 100 ng/mL), R848 (Resiquimod, 100 ng/mL) or CpG (ODN 1585, 1µM) overnight, which activated Toll-like receptor (TLR)4, TLR7/8 or TLR9 signaling, respectively. Cells were harvested for experiments. pDCs were sorted to isolate CD11c^+^B220^+^ cells using FACS cell sorter and magnetic beads (Miltenyi-Biotech, CA).

### Antibodies and flow cytometry analysis

All antibodies used for immunofluorescence staining were purchased from eBioscience (San Diego, CA), BioLegend (San Diego, CA) or BD Biosciences (San Jose, CA). Magnetic microbead-conjugated Abs and streptavidin were purchased from Miltenyi-Biotech (Auburn, CA). Recombinant human IL-2 protein was purchased from R&D Systems (Minneapolis, MN). Recombinant mouse Flt3 ligand (Flt3L) and mouse SCF were purchased from Shenandoah Biotech (Warwick, PA). Cells were stained with appropriate concentrations of mAbs. Dead cells were excluded using Fixable Viability Dye from eBioscience (San Diego, CA). Flow cytometry (FC) analyses were performed using LSRII (BD Bioscience, CA) and Canto cytometers (Becton Dickinson, NJ). Cells were sorted on BD FACSAria II (BD Bioscience, CA).

### T cell culture and activation

Splenic mononuclear cells from each group of immunized mice were cultured in the presence of DCs pulsed with or without S2P_ECTO_, multilayered S2GΔHR2-presenting E2P or I3-01v9 SApNP (1 × 10^-7^ μM) in complete IMDM medium containing IL-2 (5.0 ng/ml). Cells were collected 16 hours and 4 hours later for intracellular cytokine staining and flow cytometric analysis.

## Statistics

In antibody analysis, comparison of different vaccine groups was performed in GraphPad Prism 8.4.3 using the two-tailed unpaired Student’s *t* test. In T cell analysis, comparison of means was done using the two-tailed unpaired Student’s t test, ANOVA and then post-hoc *t* test. P values of 0.05 or less were considered significant.

## SUPPLEMENTARY MATERIALS

Supplementary material for this article is available at http://xxx/xxx/xxx.

**fig. S1**. In-vitro characterization of SARS-CoV-1/2 RBD-based immunogens.

**fig. S2**. In-vitro characterization of SARS-CoV-2 spike antigens.

**fig. S3.** In-vitro characterization of SARS-CoV-2 S2GΔHR2 SApNPs.

**fig. S4.** SARS-CoV-2 RBD/RBD-SApNP vaccine-induced binding antibody responses.

**fig. S5.** SARS-CoV-2 spike/spike-SApNP vaccine-induced binding antibody responses.

**fig. S6.** SARS-CoV-1 spike/RBD/RBD-SApNP vaccine-induced binding antibody responses.

**fig. S7.** SARS-CoV-2 RBD/RBD-SApNP vaccine-induced neutralizing antibody responses.

**fig. S8.** SARS-CoV-2 spike/spike-SApNP vaccine-induced neutralizing antibody responses.

**fig. S9.** SARS-CoV-1 spike/RBD/RBD-SApNP vaccine-induced neutralizing antibody responses.

**table. S1.** Key *in vitro* and *in vivo* characteristics of RBD and spike-based SARS-CoV-2 vaccines.

## Supporting information

Supplementary figures and ables

## Acknowledgements: Funding

This work was funded by NIH Grants AI129698 and AI140844 (to J.Z.), Ufovax/SFP-2018-0416, Ufovax/SFP-2018-1013 and Ufovax/SFP-2020-0111 (to J.Z.).

## Author contributions

Project design by L.H., Y.Z., I.A.W. and J.Z; construct design of RBD and spike-based immungoens by L.H. and J.Z.; plasmid design and processing by L.H. and C.S.; antigen production, purification and basic characterization by L.H., X.L., and T.N.; antibody production and column packing by L.H., X.L., and T.N.; antibody-antigen ELISA and BLI by L.H. and C.S.; DSC by L.H. and J.Z.; negative-stain EM by L.H. and J.Z.; plasma-antigen ELISA by L.H., X.L., and C.S.; antibody and mouse plasma neutralization by L.H. and X.L.; vaccine-induced T-cell response analysis by Y.W., C.A., and Y.Z.; Manuscript written by L.H., Y.Z., I.A.W. and J.Z. All authors were asked to comment on the manuscript. The TSRI manuscript number is 30028.

## Competing interests

Authors declare no competing interests.

## Data and materials availability

All data are available in the main text or in the supplementary materials. Additional data related to this paper may be requested from the corresponding author.

## Notes

### Competing Interest Statement

The authors have declared no competing interest.

