## Supplementary figures and ables for "Single-component, self-assembling, protein nanoparticles presenting the receptor binding domain and stabilized spike as SARS-CoV-2 vaccine candidates"

### fig. S1

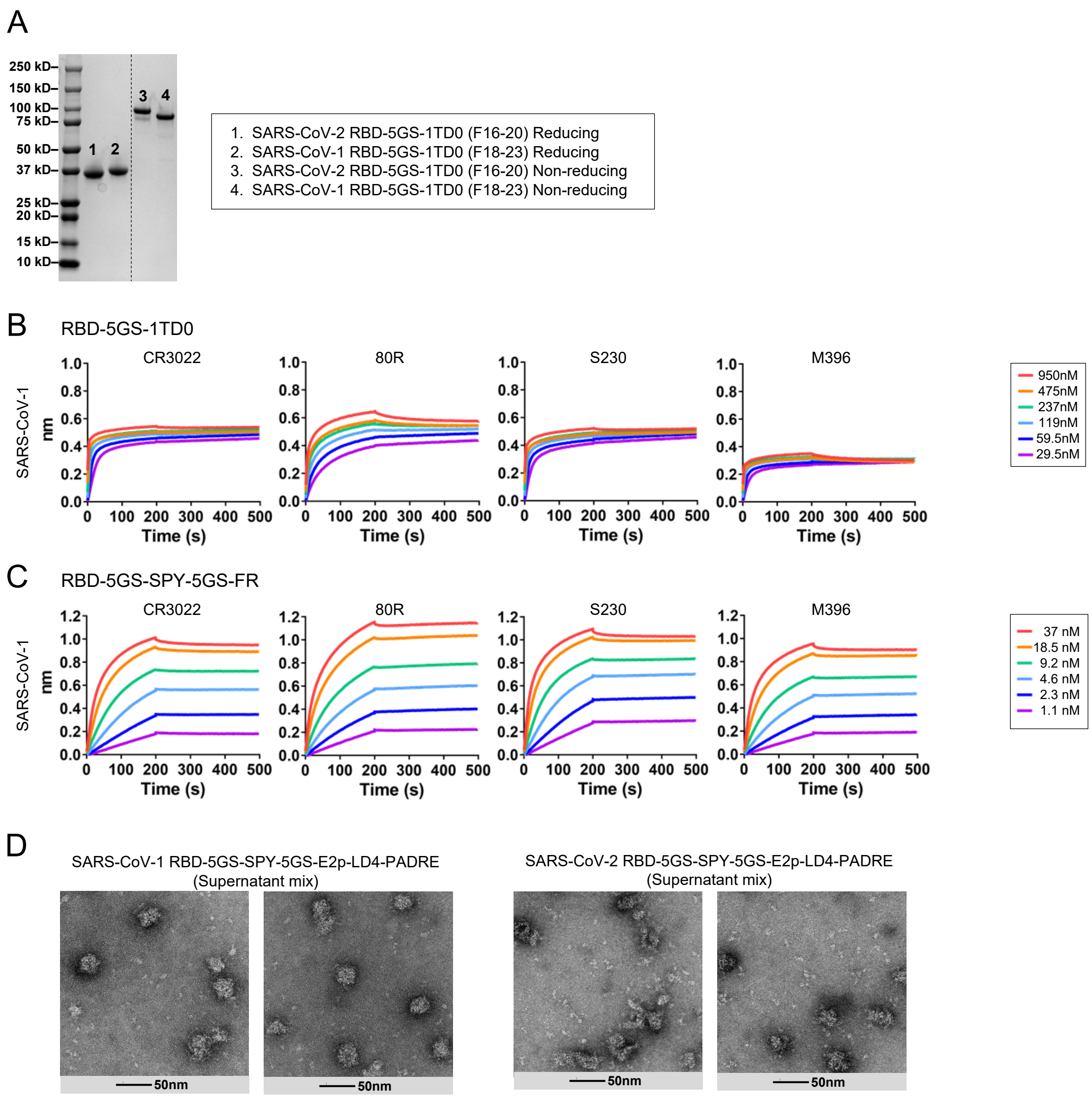


**fig. S1. In-vitro characterization of SARS-CoV-1/2 RBD-based immunogens. (A)** SDS-PAGE of SEC/CR3022-purified SARS-CoV-1/2 RBD-5GS-1TD0 under the reducing and non-reducing conditions. The black dashed line indicates where an unrelated protein analyzed on the same gel has been removed. **(B)** and **(C)** BLI profiles of SARS-CoV-1 RBD-5GS-1TD0 trimer and RBD-5GS-SPY-5GS-FR SApNP binding to four mAbs. Sensorgrams were obtained from an Octet RED96 instrument using a series of six concentrations (950-29.5 nM for the RBD trimer and 37-1.1 nM for the SPY-linked RBD FR SApNP, respectively, both by twofold dilutions) and AHC biosensors. **(D)** Negative-stain EM (nsEM) images of SARS-CoV-1/2 RBD-5GS-SPY-5GS-E2p-LD4-PADRE (or E2p-L4P) SApNPs. E2p-LD4-PADRE is a multilayered SApNP carrier with a locking domain (LD4) and a T-cell epitope (PADRE) fused to each E2p subunit. LD4 forms an inner layer to stabilize the NP shell from the inside and PADRE forms a cluster at the center of the SApNP.

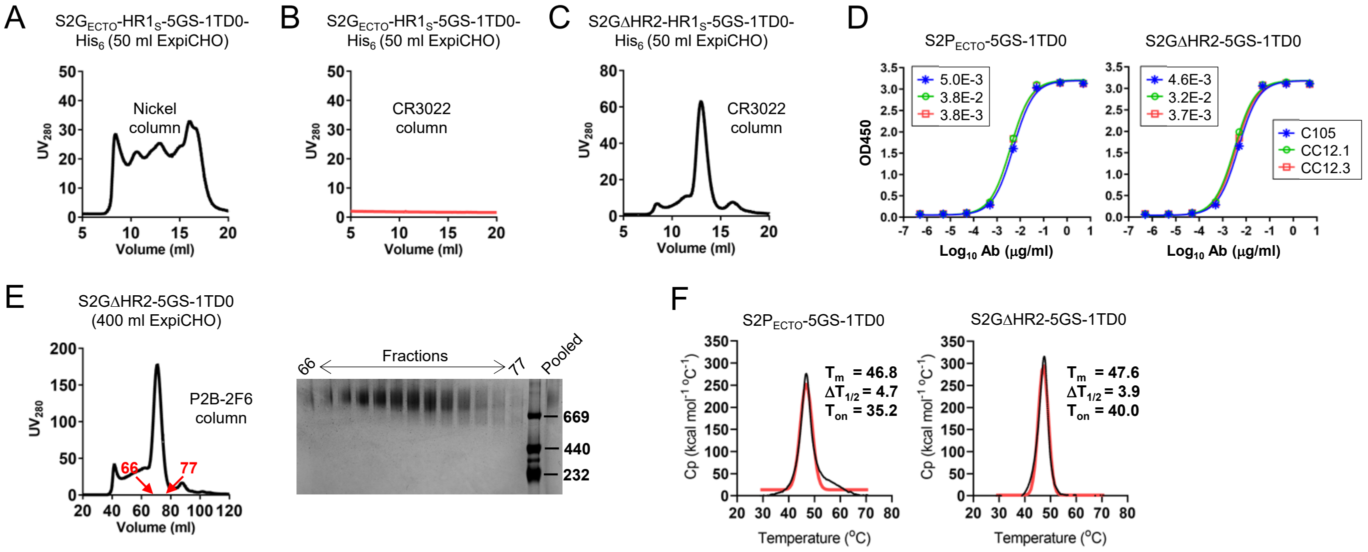


**fig. S2. In-vitro characterization of SARS-CoV-2 spike antigens. (A)** SEC profile of S2G$_{ECTO}$-HR1$_S$-5GS-1TD0-His$_6$ in which the SARS-CoV-2 HR1 region (L922-S943) is swapped with the equivalent SARS-CoV-1 HR1 region, thus termed "HR1$_S$". This construct was transiently expressed in 50 ml ExpiCHO cells and purified on a Nickel column. **(B)** SEC profile of S2G$_{ECTO}$-HR1$_S$-5GS-1TD0-His$_6$ following transient expression in 50 ml ExpiCHO cells and purification on a CR3022 column. **(C)** SEC profile of S2GΔHR2-HR1$_S$-5GS-1TD0-His$_6$ following transient expression in 50 ml ExpiCHO cells and purification on a CR3022 column. **(D)** ELISA curves of S2P$_{ECTO}$-5GS-1TD0 and S2GΔHR2-5GS-1TD0 binding to three newly identified potent human NAbs, C105, CC12.1, and CC12.3. **(E)** Left: SEC profile of S2GΔHR2-5GS-1TD0 obtained from a HiLoad 16/600 Superose 6 column following transient expression in 400 ml ExpiCHO cells and purification on a P2B-2F6 antibody column. The range of fractions used for the BN-PAGE analysis is indicated on the SEC profile. Right: BN-PAGE of SEC fractions (66-77 ml) and pooled fractions. **(F)** Thermostability of S2P$_{ECTO}$-5GS-1TD0 and S2GΔHR2-5GS-1TD0, with Tm, ΔT1/2, and Ton measured by differential scanning calorimetry (DSC) and labeled on the plots. The raw and Gaussian-fitted DSC data are shown in black and red lines, respectively.

fig. S3

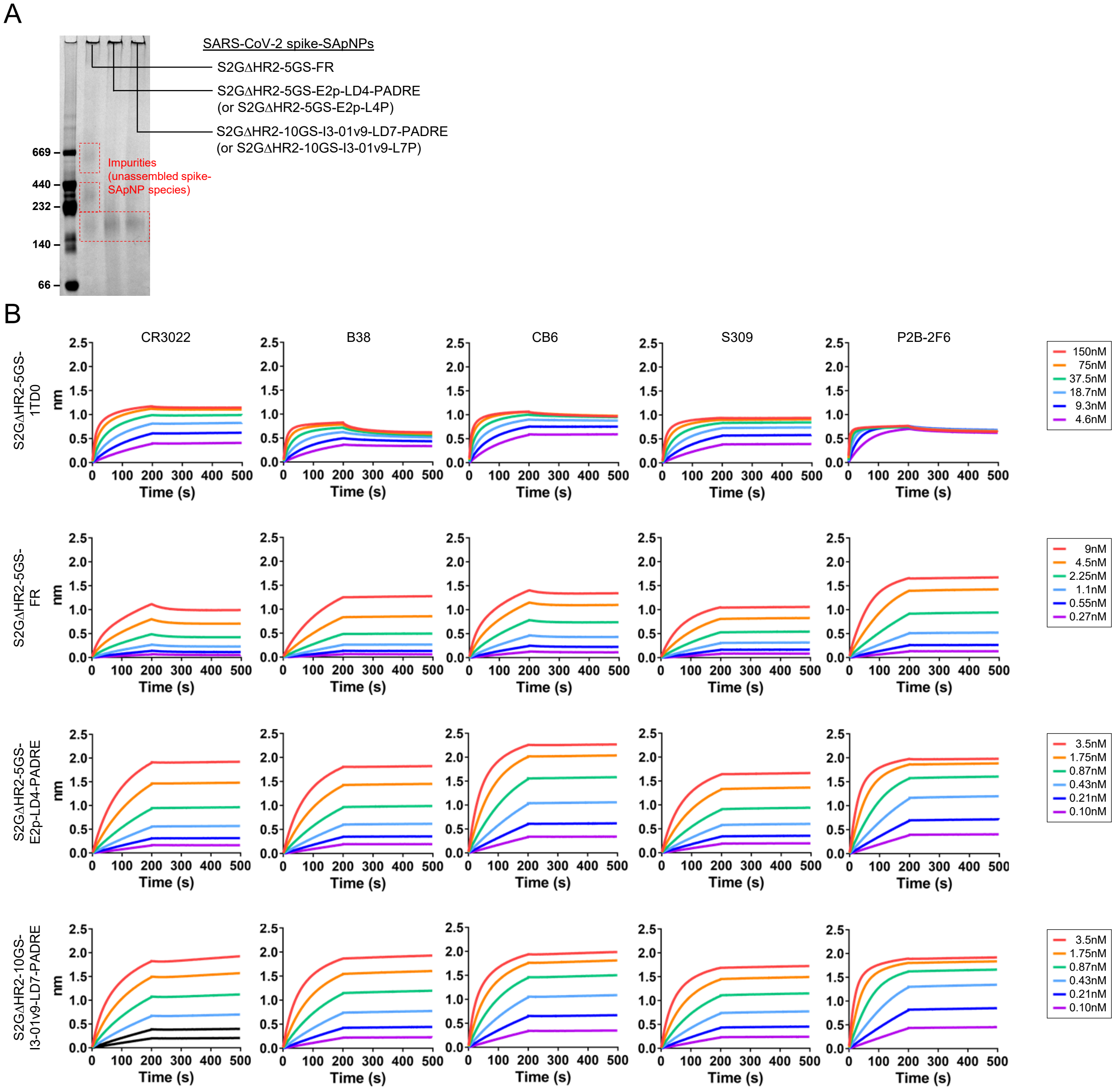


**fig. S3. In-vitro characterization of SARS-CoV-2 S2GΔHR2 SApNPs. (A)** BN-PAGE of CR3022-purified S2GΔHR2-presenting SApNPs. Due to the large size and molecular mass, NPs would be in the well on the top of the gel, whereas the unassembled species would be seen on the gel. **(B)** BLI profiles of SARS-CoV-2 S2GΔHR2-presenting SApNPs binding to five mAbs**.** The SARS-CoV-2 S2GΔHR2-5GS-1TD0 trimer was included for comparison. Sensorgrams were obtained from an Octet RED96 instrument using a series of six concentrations (150-4.6nM for the S2GΔHR2 trimer, 9-0.27nM for the FR NP, and 3.5-0.1nM for the E2p and I3-01v9 NPs, respectively, all by twofold dilutions) and AHQ biosensors.

#### fig. S4

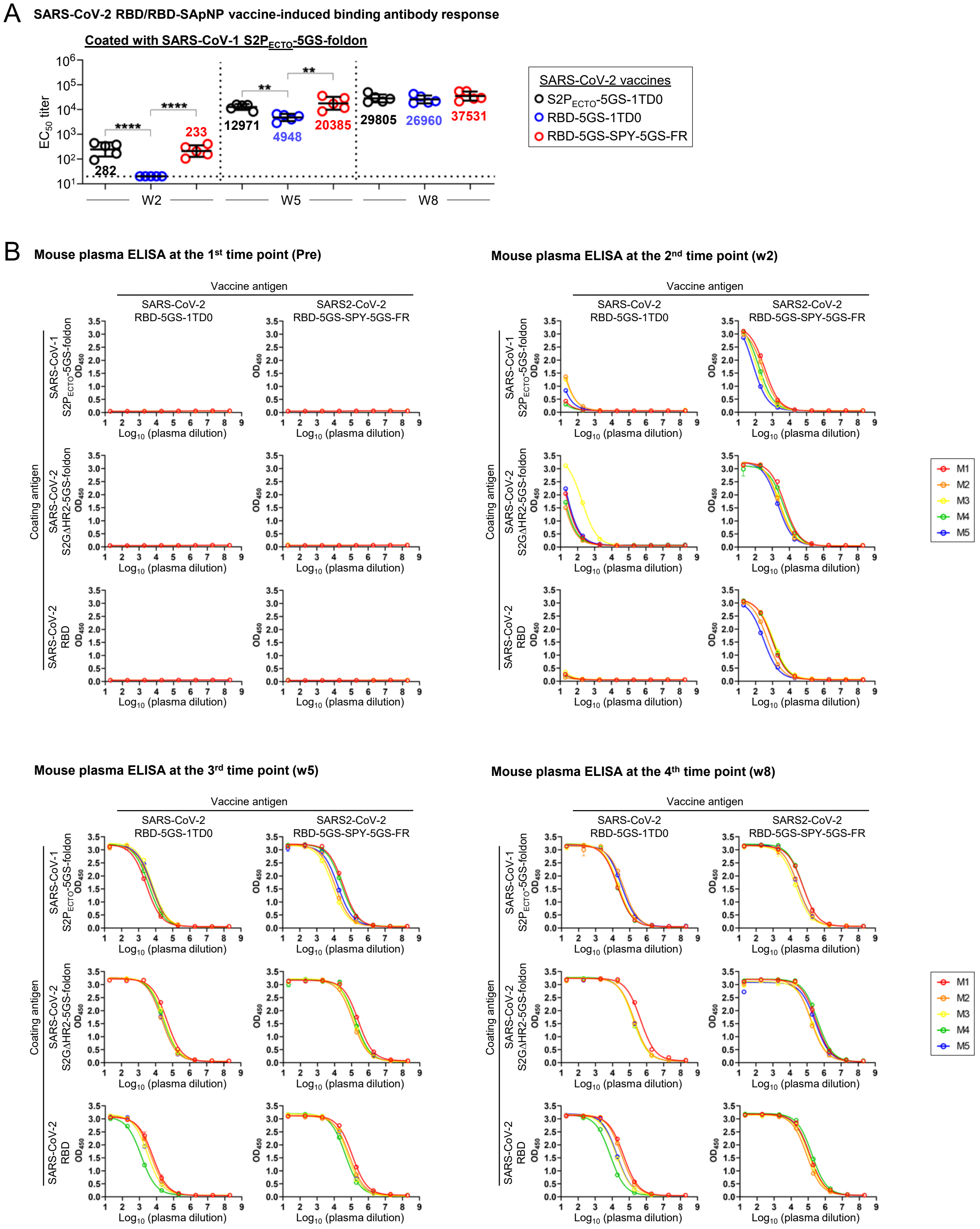

A
SARS-CoV-2 RBD/RBD-SApNP vaccine-induced binding antibody response
Coated with SARS-CoV-1 S2P$_{ECTO}$-5GS-foldon
EC$_{50}$ titer
****
****
282
233
**
**
12971
4948
20385
29805
26960
37531
W2
W5
W8
SARS-CoV-2 vaccines
S2P$_{ECTO}$-5GS-1TD0
RBD-5GS-1TD0
RBD-5GS-SPY-5GS-FR
B
Mouse plasma ELISA at the 1$^{st}$ time point (Pre)
Mouse plasma ELISA at the 2$^{nd}$ time point (w2)
Mouse plasma ELISA at the 3$^{rd}$ time point (w5)
Mouse plasma ELISA at the 4$^{th}$ time point (w8)
Vaccine antigen
SARS-CoV-2 RBD-5GS-1TD0
SARS2-CoV-2 RBD-5GS-SPY-5GS-FR
Coating antigen
SARS-CoV-1 S2P$_{ECTO}$-5GS-foldon
SARS-CoV-2 S2GΔHR2-5GS-foldon
SARS-CoV-2 RBD
OD$_{450}$
Log$_{10}$ (plasma dilution)
M1
M2
M3
M4
M5

### fig. S4

**Mouse plasma ELISA at the 5th time point (w11)**

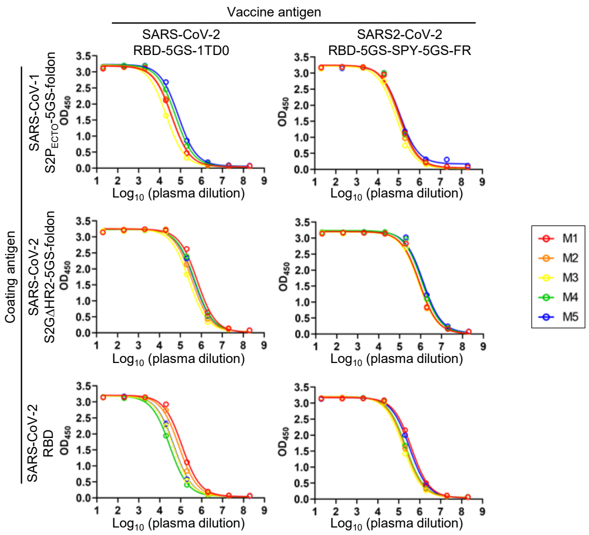


C **Mouse plasma ELISA $EC_{50}$ titers**

Coating antigen: SARS-CoV-1 S2P$_{ECTO}$-5GS-foldon

| SARS-CoV-2 vaccine antigen | w2 | | | | | w5 | | | | | w8 | | | | | w11 | | | | |
|---|---|---|---|---|---|---|---|---|---|---|---|---|---|---|---|---|---|---|---|---|
| | M1 | M2 | M3 | M4 | M5 | M1 | M2 | M3 | M4 | M5 | M1 | M2 | M3 | M4 | M5 | M1 | M2 | M3 | M4 | M5 |
| RBD-5GS-1TD0 | <20 | <20 | <20 | <20 | <20 | 2849 | 5426 | 6544 | 4016 | 5908 | 18964 | 42380 | 21799 | 18353 | 33308 | 37799 | 38906 | 21152 | 57386 | 77425 |
| RBD-5GS-SPY-5GS-FR | 402 | 299 | 155 | 205 | 105 | 35702 | 11730 | 8478 | 28869 | 17148 | 55019 | 29258 | 21429 | 54316 | 27631 | 116637 | 100384 | 75563 | 114456 | 117834 |

Coating antigen: SARS-CoV-2 S2GΔHR2-5GS-foldon

| SARS-CoV-2 vaccine antigen | w2 | | | | | w5 | | | | | w8 | | | | | w11 | | | | |
|---|---|---|---|---|---|---|---|---|---|---|---|---|---|---|---|---|---|---|---|---|
| | M1 | M2 | M3 | M4 | M5 | M1 | M2 | M3 | M4 | M5 | M1 | M2 | M3 | M4 | M5 | M1 | M2 | M3 | M4 | M5 |
| RBD-5GS-1TD0 | 33 | <20 | 237 | 23 | 41 | 45821 | 22822 | 31754 | 28895 | 23447 | 437779 | 186022 | 163880 | 159780 | 177684 | 640237 | 365517 | 258393 | 495393 | 458394 |
| RBD-5GS-SPY-5GS-FR | 5719 | 2893 | 3239 | 4472 | 2280 | 245016 | 119447 | 145979 | 172388 | 146375 | 343504 | 169804 | 194569 | 415264 | 305726 | 893358 | 866269 | 801461 | 1181201 | 1352045 |

Coating antigen: SARS-CoV-2 RBD

| SARS-CoV-2 vaccine antigen | w2 | | | | | w5 | | | | | w8 | | | | | w11 | | | | |
|---|---|---|---|---|---|---|---|---|---|---|---|---|---|---|---|---|---|---|---|---|
| | M1 | M2 | M3 | M4 | M5 | M1 | M2 | M3 | M4 | M5 | M1 | M2 | M3 | M4 | M5 | M1 | M2 | M3 | M4 | M5 |
| RBD-5GS-1TD0 | <20 | <20 | <20 | <20 | <20 | 6061 | 4770 | 3127 | 1293 | 3140 | 46119 | 35578 | 17646 | 7671 | 18814 | 116486 | 82504 | 45221 | 30347 | 47738 |
| RBD-5GS-SPY-5GS-FR | 913 | 499 | 1092 | 978 | 292 | 115261 | 78765 | 61890 | 45215 | 61522 | 125683 | 81629 | 110472 | 165129 | 110446 | 402781 | 200112 | 165137 | 217751 | 328643 |

**fig. S4. SARS-CoV-2 RBD/RBD-SApNP vaccine-induced binding antibody responses. (A)** $EC_{50}$ titers derived from the ELISA binding of mouse plasma from two SARS-CoV-2 RBD-based vaccine groups (RBD-5GS-1TD0 and RBD-5GS-SPY-5GS-FR) to the coating antigen (SARS-CoV-1 S2P$_{ECTO}$-5GS-foldon) are plotted, with average $EC_{50}$ values labeled on the plots. The S2P spike vaccine group (S2P$_{ECTO}$-5GS-1TD0) is included here for comparison with RBD-based vaccine groups, with the detailed ELISA data shown in fig. S5. The *P*-values were determined by an unpaired *t* test in GraphPad Prism 8.4.3 with (*) indicating the level of statistical significance (*: $0.01<P\leq0.05$; **: $0.001<P\leq0.01$; ***: $0.0001<P\leq0.001$; ****: $P\leq0.0001$). **(B)** ELISA binding curves of mouse plasma from two SARS-CoV-2 RBD-based vaccine groups to three coating antigens, SARS-CoV-1 S2P$_{ECTO}$-5GS-foldon, SARS-CoV-2 S2GΔHR2-5GS-foldon, and SARS-CoV-2 RBD. **(C)** Summary of $EC_{50}$ titers measured for two SARS-CoV-2 RBD-based vaccine groups against three coating antigens. Color coding indicates the level of $EC_{50}$ titer (white: no binding; green to red: low to high). The $EC_{50}$ values were calculated in GraphPad Prism 8.4.3. Of note, the $EC_{50}$ values at w2 were derived by setting the lower/upper constraints of $OD_{450}$ at 0.0/3.2 to achieve greater accuracy.

**fig. S5**

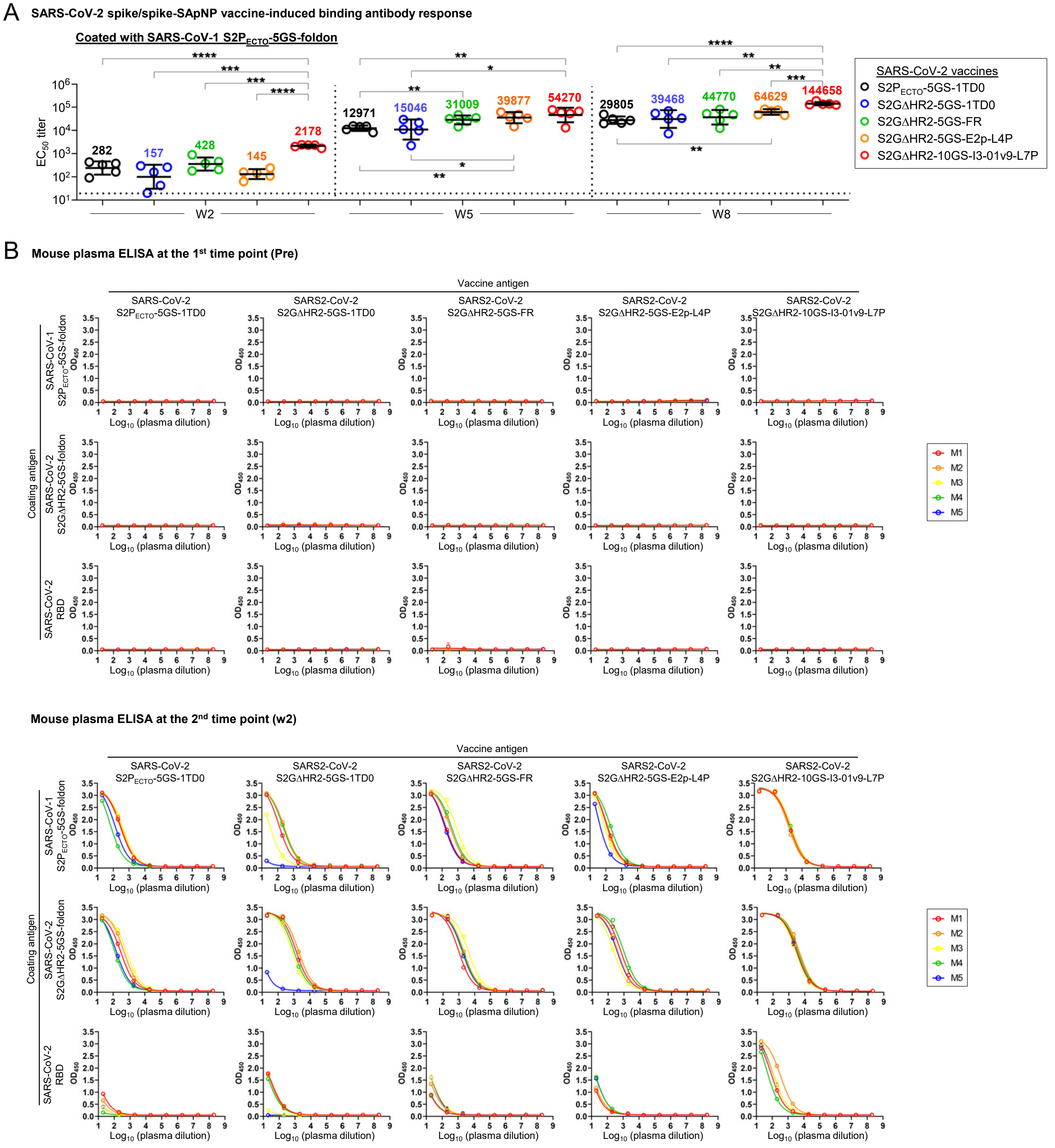
A
SARS-CoV-2 spike/spike-SApNP vaccine-induced binding antibody response
Coated with SARS-CoV-1 S2P$_{ECTO}$-5GS-foldon
EC$_{50}$ titer
282
157
428
145
2178
12971
15046
31009
39877
54270
29805
39468
44770
64629
144658
W2
W5
W8
SARS-CoV-2 vaccines
S2P$_{ECTO}$-5GS-1TD0
S2GΔHR2-5GS-1TD0
S2GΔHR2-5GS-FR
S2GΔHR2-5GS-E2p-L4P
S2GΔHR2-10GS-I3-01v9-L7P
B
Mouse plasma ELISA at the 1st time point (Pre)
Vaccine antigen
SARS-CoV-2 S2P$_{ECTO}$-5GS-1TD0
SARS2-CoV-2 S2GΔHR2-5GS-1TD0
SARS2-CoV-2 S2GΔHR2-5GS-FR
SARS2-CoV-2 S2GΔHR2-5GS-E2p-L4P
SARS2-CoV-2 S2GΔHR2-10GS-I3-01v9-L7P
Coating antigen
SARS-CoV-1 S2P$_{ECTO}$-5GS-foldon
SARS-CoV-2 S2GΔHR2-5GS-foldon
SARS-CoV-2 RBD
OD$_{450}$
Log$_{10}$ (plasma dilution)
M1
M2
M3
M4
M5
Mouse plasma ELISA at the 2nd time point (w2)
Vaccine antigen
SARS-CoV-2 S2P$_{ECTO}$-5GS-1TD0
SARS2-CoV-2 S2GΔHR2-5GS-1TD0
SARS2-CoV-2 S2GΔHR2-5GS-FR
SARS2-CoV-2 S2GΔHR2-5GS-E2p-L4P
SARS2-CoV-2 S2GΔHR2-10GS-I3-01v9-L7P
Coating antigen
SARS-CoV-1 S2P$_{ECTO}$-5GS-foldon
SARS-CoV-2 S2GΔHR2-5GS-foldon
SARS-CoV-2 RBD
OD$_{450}$
Log$_{10}$ (plasma dilution)
M1
M2
M3
M4
M5

**fig. S5**

**Mouse plasma ELISA at the 3rd time point (w5)**

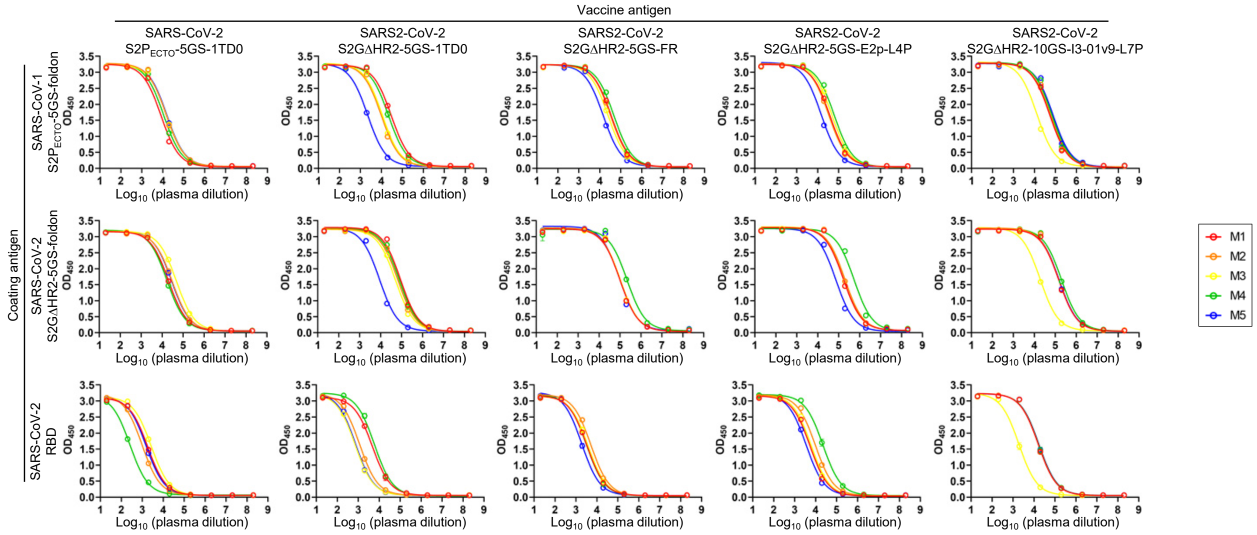


**Mouse plasma ELISA at the 4th time point (w8)**

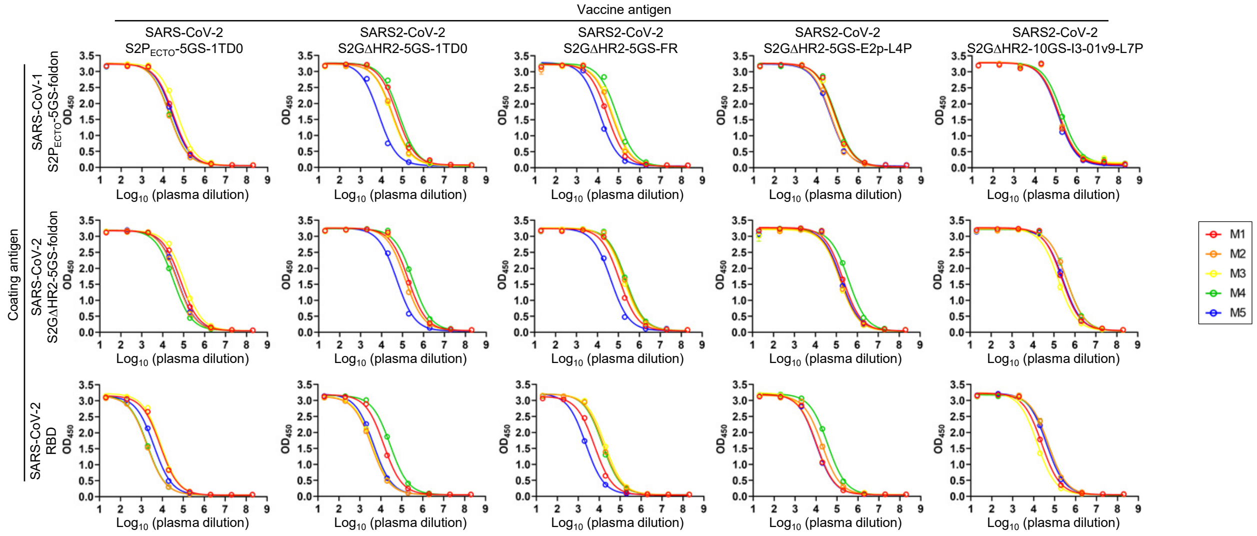

**Mouse plasma ELISA at the 5th time point (w11)**

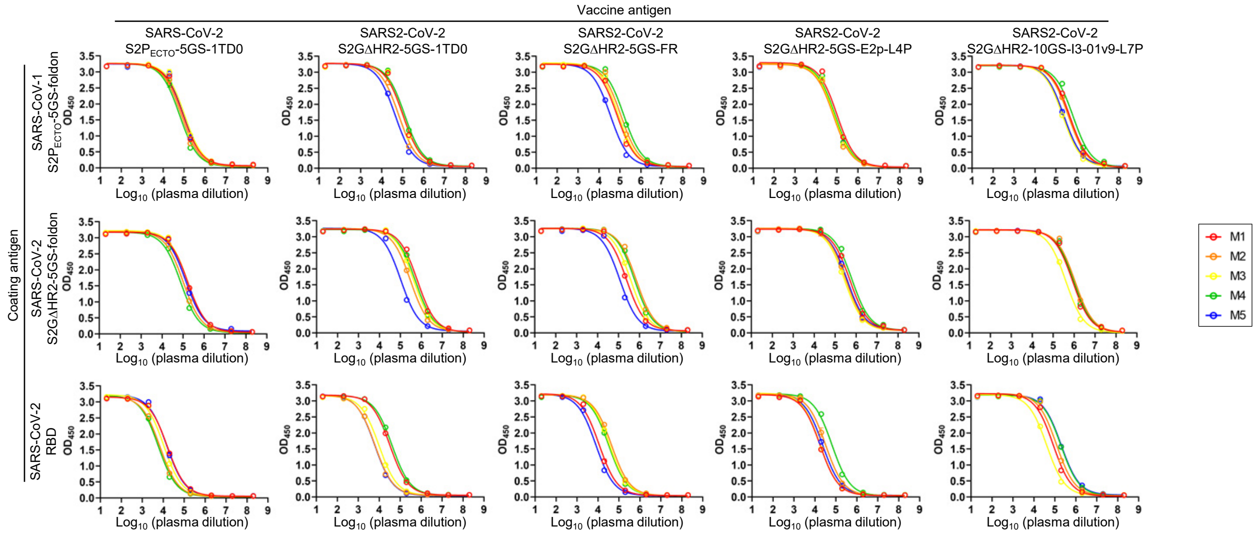


C **Mouse plasma ELISA $EC_{50}$ titers**

Coating antigen: SARS-CoV-1 $S2P_{ECTO}$-5GS-foldon

| SARS-CoV-2 vaccine antigen | w2 | | | | | w5 | | | | | w8 | | | | | w11 | | | | |
|---|---|---|---|---|---|---|---|---|---|---|---|---|---|---|---|---|---|---|---|---|
| | M1 | M2 | M3 | M4 | M5 | M1 | M2 | M3 | M4 | M5 | M1 | M2 | M3 | M4 | M5 | M1 | M2 | M3 | M4 | M5 |
| $S2P_{ECTO}$-5GS-1TD0 | 337 | 378 | 438 | 92 | 166 | 8229 | 14598 | 15271 | 11029 | 15730 | 31346 | 19923 | 49495 | 20899 | 27362 | 87963 | 74675 | 105473 | 61814 | 98729 |
| S2GDHR2-5GS-1TD0 | 160 | 300 | 43 | 264 | <20 | 29889 | 9934 | 11211 | 21972 | 2225 | 54736 | 33977 | 31597 | 69701 | 7331 | 104856 | 65140 | 102307 | 125912 | 44213 |
| S2GDHR2-5GS-FR | 207 | 379 | 907 | 463 | 182 | 35236 | 32621 | 25025 | 47534 | 14631 | 28314 | 48030 | 53004 | 81962 | 12542 | 73512 | 110204 | 85157 | 152546 | 35108 |
| S2GDHR2-5GS-E2p-L4P | 161 | 145 | 111 | 244 | 64 | 38631 | 34068 | 49277 | 62620 | 14788 | 76610 | 48384 | 69459 | 83674 | 45019 | 101670 | 68765 | 75147 | 83008 | 78758 |
| S2GDHR2-10GS-I3-01v9-L7P | 2047 | 1697 | 2368 | 2434 | 2342 | 53863 | 59267 | 13164 | 67367 | 77688 | 136410 | 139928 | 130590 | 190016 | 126344 | 461522 | 400892 | 212809 | 662353 | 240540 |

Coating antigen: SARS-CoV-2 S2GΔHR2-5GS-foldon

| SARS-CoV-2 vaccine antigen | w2 | | | | | w5 | | | | | w8 | | | | | w11 | | | | |
|---|---|---|---|---|---|---|---|---|---|---|---|---|---|---|---|---|---|---|---|---|
| | M1 | M2 | M3 | M4 | M5 | M1 | M2 | M3 | M4 | M5 | M1 | M2 | M3 | M4 | M5 | M1 | M2 | M3 | M4 | M5 |
| $S2P_{ECTO}$-5GS-1TD0 | 342 | 564 | 786 | 152 | 186 | 20525 | 30418 | 47919 | 16989 | 28307 | 71336 | 48544 | 104033 | 32473 | 51430 | 168104 | 101372 | 164947 | 77599 | 136078 |
| S2GDHR2-5GS-1TD0 | 1623 | 2329 | 915 | 1190 | <20 | 88704 | 66897 | 51587 | 77015 | 8882 | 188541 | 133257 | 137492 | 308124 | 53005 | 648885 | 297906 | 427211 | 524920 | 103434 |
| S2GDHR2-5GS-FR | 1168 | 2365 | 4609 | 2102 | 1925 | 100424 | 99021 | 105640 | 217243 | 94779 | 112191 | 207821 | 177619 | 227282 | 43471 | 240083 | 710983 | 381026 | 592260 | 116127 |
| S2GDHR2-5GS-E2p-L4P | 748 | 481 | 268 | 1132 | 435 | 168389 | 197689 | 167303 | 549503 | 74194 | 209083 | 153974 | 150854 | 389845 | 173525 | 500160 | 324937 | 276279 | 631496 | 369062 |
| S2GDHR2-10GS-I3-01v9-L7P | 3695 | 5177 | 3135 | 4338 | 3353 | 147140 | 146498 | 19018 | 197789 | 152653 | 239618 | 402217 | 162393 | 412328 | 272905 | 843216 | 1100695 | 403762 | 949208 | 944347 |

Coating antigen: SARS-CoV-2 RBD

| SARS-CoV-2 vaccine antigen | w2 | | | | | w5 | | | | | w8 | | | | | w11 | | | | |
|---|---|---|---|---|---|---|---|---|---|---|---|---|---|---|---|---|---|---|---|---|
| | M1 | M2 | M3 | M4 | M5 | M1 | M2 | M3 | M4 | M5 | M1 | M2 | M3 | M4 | M5 | M1 | M2 | M3 | M4 | M5 |
| $S2P_{ECTO}$-5GS-1TD0 | <20 | <20 | <20 | <20 | <20 | 1801 | 980 | 2693 | 260 | 1533 | 7881 | 1861 | 8448 | 1952 | 3948 | 15828 | 6935 | 10668 | 5826 | 14807 |
| S2GDHR2-5GS-1TD0 | 26 | 24 | <20 | <20 | <20 | 4693 | 1161 | 703 | 5881 | 751 | 13835 | 3578 | 3906 | 28190 | 4493 | 32268 | 6328 | 9751 | 40928 | 6118 |
| S2GDHR2-5GS-FR | <20 | <20 | 22 | <20 | 21 | 3340 | 5205 | 3799 | 3229 | 1900 | 5996 | 15596 | 18918 | 14222 | 2293 | 11360 | 44589 | 35885 | 32606 | 7947 |
| S2GDHR2-5GS-E2p-L4P | <20 | <20 | <20 | 21 | <20 | 5425 | 9801 | 6186 | 22466 | 3554 | 10682 | 19414 | 10367 | 36238 | 10091 | 17641 | 32324 | 17662 | 64941 | 23122 |
| S2GDHR2-10GS-I3-01v9-L7P | 112 | 384 | 159 | 73 | 151 | 16265 | 16054 | 1920 | 16797 | 17368 | 24415 | 49820 | 15026 | 51700 | 41325 | 84065 | 120318 | 40866 | 194224 | 199672 |

**fig. S5. SARS-CoV-2 spike/spike-SApNP vaccine-induced binding antibody responses. (A)** $EC_{50}$ titers derived from the ELISA binding of mouse plasma from five SARS-CoV-2 spike-based vaccine groups ($S2P_{ECTO}$-5GS-1TD0, S2GΔHR2-5GS-1TD0, S2GΔHR2-5GS-FR, S2GΔHR2-5GS-E2p-L4P, and S2GΔHR2-10GS-I3-01v9-L7P) to the coating antigen (SARS-CoV-1 S2P-5GS-foldon) are plotted, with average $EC_{50}$ values labeled on the plots. The S2P spike vaccine group ($S2P_{ECTO}$-5GS-1TD0) is included here for comparison with S2GΔHR2 spike-based vaccine groups. The *P*-values were determined by an unpaired *t* test in GraphPad Prism 8.4.3 with (*) indicating the level of statistical significance (*: $0.01<P\leq0.05$; **: $0.001<P\leq0.01$; ***: $0.0001<P\leq0.001$; ****: $P\leq0.0001$). **(B)** ELISA binding curves of mouse plasma from five SARS-CoV-2 spike-based vaccine groups to three coating antigens, SARS-CoV-1 $S2P_{ECTO}$-5GS-foldon, SARS-CoV-2 S2GΔHR2-5GS-foldon, and SARS-CoV-2 RBD. **(C)** Summary of $EC_{50}$ titers measured for five SARS-CoV-2 spike-based vaccine groups against three coating antigens. Color coding indicates the level of $EC_{50}$ titer (white: no binding; green to red: low to high). The $EC_{50}$ values were calculated in GraphPad Prism 8.4.3. Of note, the $EC_{50}$ values at w2 were derived by setting the lower/upper constraints of $OD_{450}$ at 0.0/3.2 to achieve greater accuracy.

### fig. S6

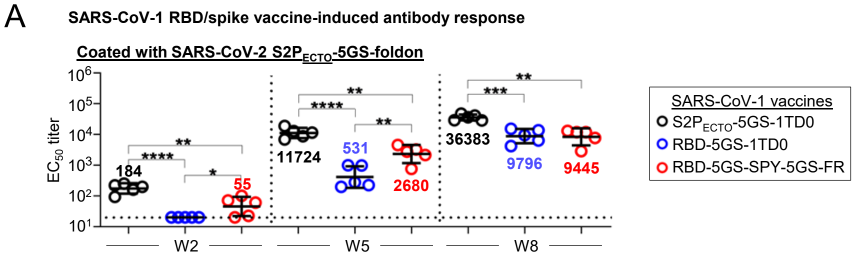

A
SARS-CoV-1 RBD/spike vaccine-induced antibody response
Coated with SARS-CoV-2 S2PECTO-5GS-foldon
EC50 titer
184
55
11724
531
2680
36383
9796
9445
W2
W5
W8
SARS-CoV-1 vaccines
S2PECTO-5GS-1TD0
RBD-5GS-1TD0
RBD-5GS-SPY-5GS-FR


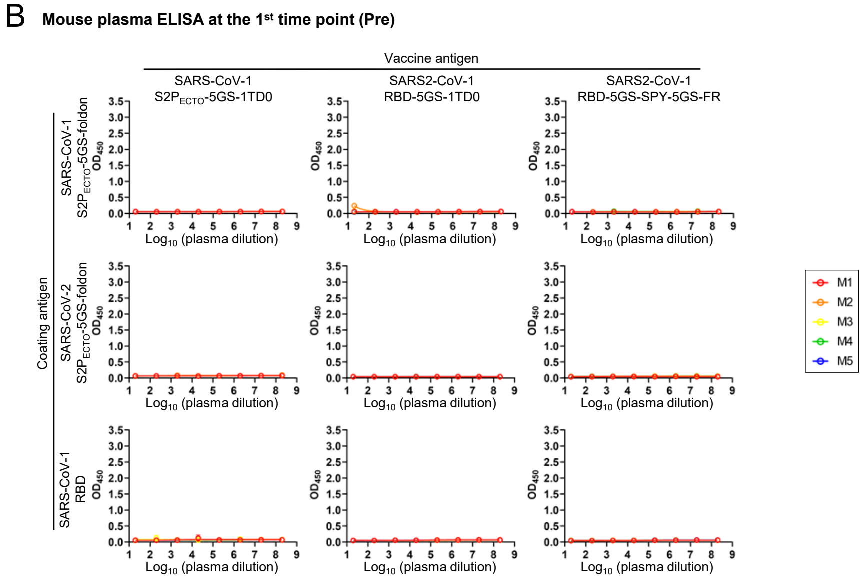

B
Mouse plasma ELISA at the 1st time point (Pre)
Vaccine antigen
SARS-CoV-1 S2PECTO-5GS-1TD0
SARS2-CoV-1 RBD-5GS-1TD0
SARS2-CoV-1 RBD-5GS-SPY-5GS-FR
Coating antigen
SARS-CoV-1 S2PECTO-5GS-foldon
SARS-CoV-2 S2PECTO-5GS-foldon
SARS-CoV-1 RBD
OD450
Log10 (plasma dilution)
M1
M2
M3
M4
M5


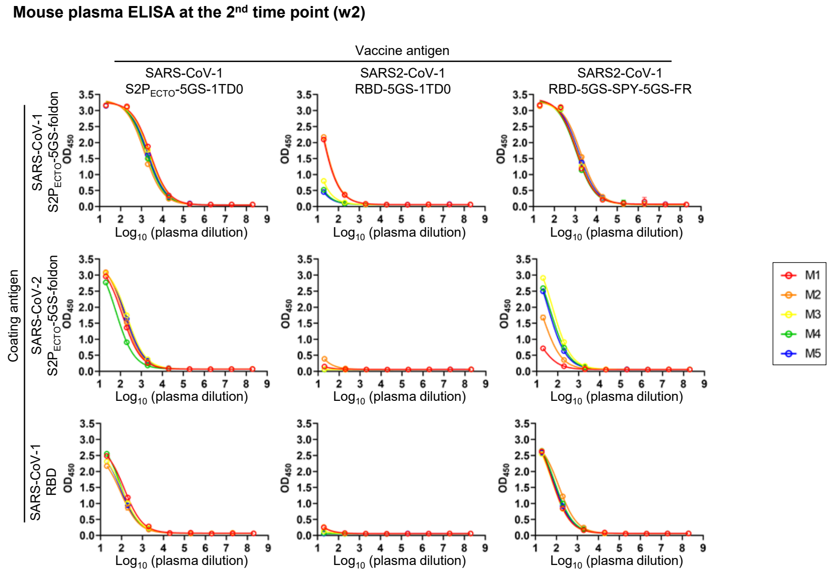

Mouse plasma ELISA at the 2nd time point (w2)
Vaccine antigen
SARS-CoV-1 S2PECTO-5GS-1TD0
SARS2-CoV-1 RBD-5GS-1TD0
SARS2-CoV-1 RBD-5GS-SPY-5GS-FR
Coating antigen
SARS-CoV-1 S2PECTO-5GS-foldon
SARS-CoV-2 S2PECTO-5GS-foldon
SARS-CoV-1 RBD
OD450
Log10 (plasma dilution)
M1
M2
M3
M4
M5

**fig. S6**

**Mouse plasma ELISA at the 3rd time point (w5)**

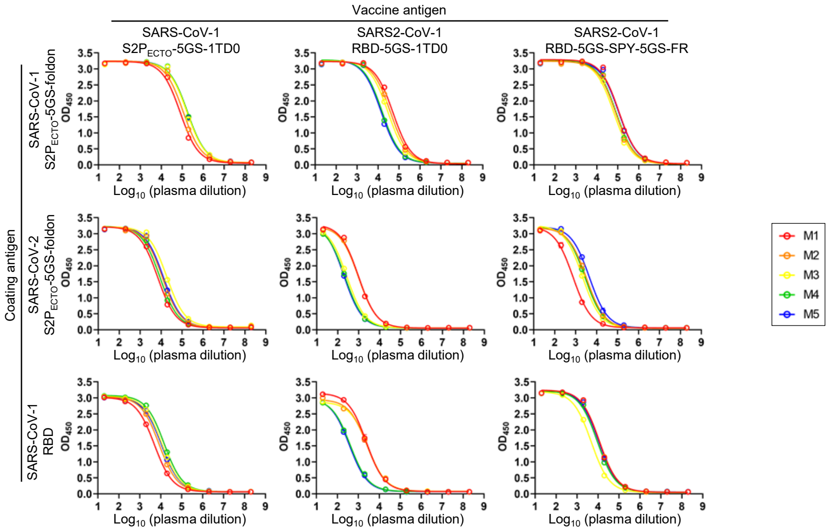


**Mouse plasma ELISA at the 4th time point (w8)**

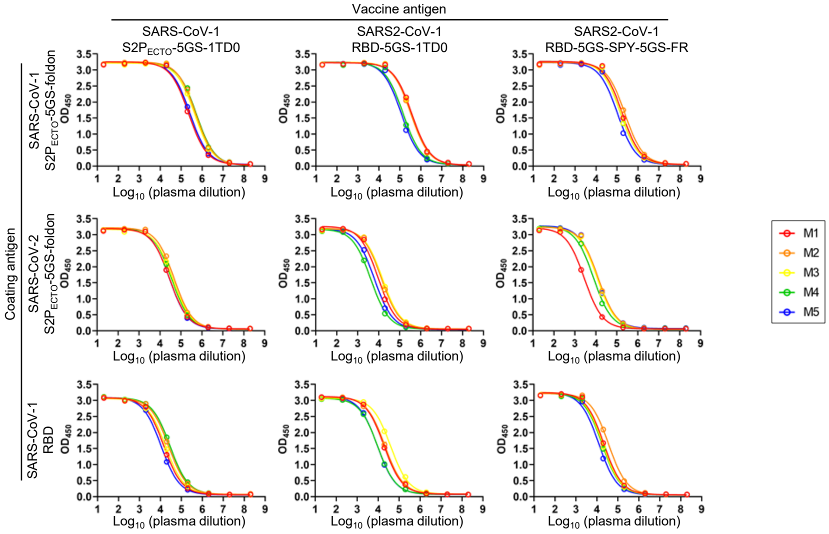

### fig. S6

**Mouse plasma ELISA at the 5th time point (w11)**

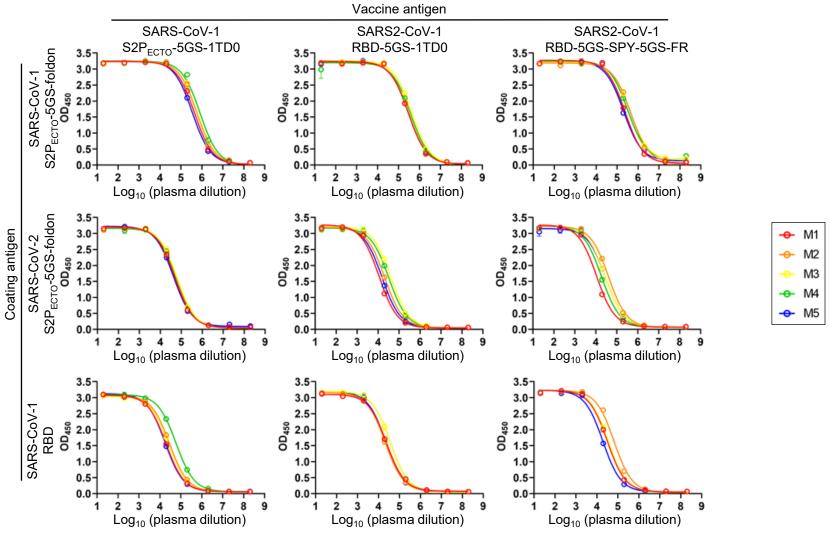


C **Mouse plasma ELISA $EC_{50}$ titers**

Coating antigen: SARS-CoV-1 S2P$_{ECTO}$-5GS-foldon

| SARS-CoV-1 vaccine antigen | w2 M1 | w2 M2 | w2 M3 | w2 M4 | w2 M5 | w5 M1 | w5 M2 | w5 M3 | w5 M4 | w5 M5 | w8 M1 | w8 M2 | w8 M3 | w8 M4 | w8 M5 | w11 M1 | w11 M2 | w11 M3 | w11 M4 | w11 M5 |
|---|---|---|---|---|---|---|---|---|---|---|---|---|---|---|---|---|---|---|---|---|
| S2P$_{ECTO}$-5GS-1TD0 | 2874 | 1611 | 2546 | 1924 | 2196 | 79871 | 111383 | 167209 | 179036 | 170765 | 226510 | 499160 | 444229 | 539886 | 264941 | 449939 | 577309 | 607415 | 818477 | 352607 |
| RBD-5GS-1TD0 | 35 | 38 | <20 | <20 | <20 | 51448 | 39666 | 29849 | 15658 | 13955 | 378373 | 337915 | 353896 | 144261 | 117614 | 290621 | 294825 | 425889 | 370917 | 290601 |
| RBD-5GS-SPY-5GS-FR | 1410 | 2047 | 1496 | 1320 | 1692 | 113510 | 78891 | 64381 | 83234 | 111601 | 178346 | 237355 | 151022 | 147743 | 107270 | 242440 | 454498 | 412469 | 328237 | 204991 |

Coating antigen: SARS-CoV-2 S2P$_{ECTO}$-5GS-foldon

| SARS-CoV-1 vaccine antigen | w2 M1 | w2 M2 | w2 M3 | w2 M4 | w2 M5 | w5 M1 | w5 M2 | w5 M3 | w5 M4 | w5 M5 | w8 M1 | w8 M2 | w8 M3 | w8 M4 | w8 M5 | w11 M1 | w11 M2 | w11 M3 | w11 M4 | w11 M5 |
|---|---|---|---|---|---|---|---|---|---|---|---|---|---|---|---|---|---|---|---|---|
| S2P$_{ECTO}$-5GS-1TD0 | 159 | 196 | 251 | 92 | 222 | 6913 | 11383 | 18692 | 8754 | 12877 | 28975 | 47458 | 41115 | 36059 | 28310 | 49277 | 52192 | 62700 | 60806 | 42133 |
| RBD-5GS-1TD0 | <20 | <20 | <20 | <20 | <20 | 985 | 975 | 274 | 224 | 197 | 9476 | 13677 | 15302 | 4216 | 6307 | 11223 | 20517 | 40915 | 32465 | 15295 |
| RBD-5GS-SPY-5GS-FR | <20 | 22 | 100 | 70 | 60 | 751 | 3243 | 2124 | 2768 | 4513 | 2840 | 13086 | 11753 | 7693 | 11851 | 10886 | 40615 | 28253 | 18689 | 29739 |

Coating antigen: SARS-CoV-1 RBD

| SARS-CoV-1 vaccine antigen | w2 M1 | w2 M2 | w2 M3 | w2 M4 | w2 M5 | w5 M1 | w5 M2 | w5 M3 | w5 M4 | w5 M5 | w8 M1 | w8 M2 | w8 M3 | w8 M4 | w8 M5 | w11 M1 | w11 M2 | w11 M3 | w11 M4 | w11 M5 |
|---|---|---|---|---|---|---|---|---|---|---|---|---|---|---|---|---|---|---|---|---|
| S2$_{PECTO}$-5GS-1TD0 | 104 | 55 | 73 | 89 | 68 | 4967 | 8355 | 11547 | 14466 | 10683 | 14673 | 24576 | 17828 | 28878 | 10869 | 19455 | 28689 | 26660 | 58793 | 17679 |
| RBD-5GS-1TD0 | <20 | <20 | <20 | <20 | <20 | 2446 | 2600 | 2827 | 415 | 371 | 18820 | 20963 | 39469 | 9503 | 9100 | 24044 | 20641 | 35559 | 36058 | 22822 |
| RBD-5GS-SPY-5GS-FR | 79 | 118 | 74 | 94 | 83 | 11483 | 11758 | 4772 | 9294 | 10888 | 21284 | 36104 | 18587 | 17645 | 12351 | 31380 | 66025 | 34973 | 34373 | 19241 |

**fig. S6. SARS-CoV-1 spike/RBD/RBD-SApNP vaccine-induced binding antibody responses. (A)** $EC_{50}$ titers derived from the ELISA binding of mouse plasma from three SARS-CoV-1 vaccine groups (S2P$_{ECTO}$-5GS-1TD0, RBD-5GS-1TD0, and RBD-5GS-SPY-5GS-FR) to the coating antigen (SARS-CoV-2 S2P$_{ECTO}$-5GS-foldon) are plotted, with average $EC_{50}$ values labeled on the plots. The S2P spike vaccine group (S2P$_{ECTO}$-5GS-1TD0) is included here for comparison with RBD-based vaccine groups. The *P*-values were determined by an unpaired *t* test in GraphPad Prism 8.4.3 with (*) indicating the level of statistical significance (*: $0.01<P\leq0.05$; **: $0.001<P\leq0.01$; ***: $0.0001<P\leq0.001$; ****: $P\leq0.0001$). **(B)** ELISA binding curves of mouse plasma from three SARS-CoV-1 vaccine groups to three coating antigens, SARS-CoV-1 S2P-5GS-foldon, SARS-CoV-2 S2P$_{ECTO}$-5GS-foldon, and SARS-CoV-1 RBD. **(C)** Summary of $EC_{50}$ titers measured for three SARS-CoV-1 vaccine groups against three coating antigens. Color coding indicates the level of $EC_{50}$ titer (white: no binding; green to red: low to high). The $EC_{50}$ values were calculated in GraphPad Prism 8.4.3. Of note, the $EC_{50}$ values at w2 were derived by setting the lower/upper constraints of $OD_{450}$ at 0.0/3.2 to achieve greater accuracy.

### fig. S7

A

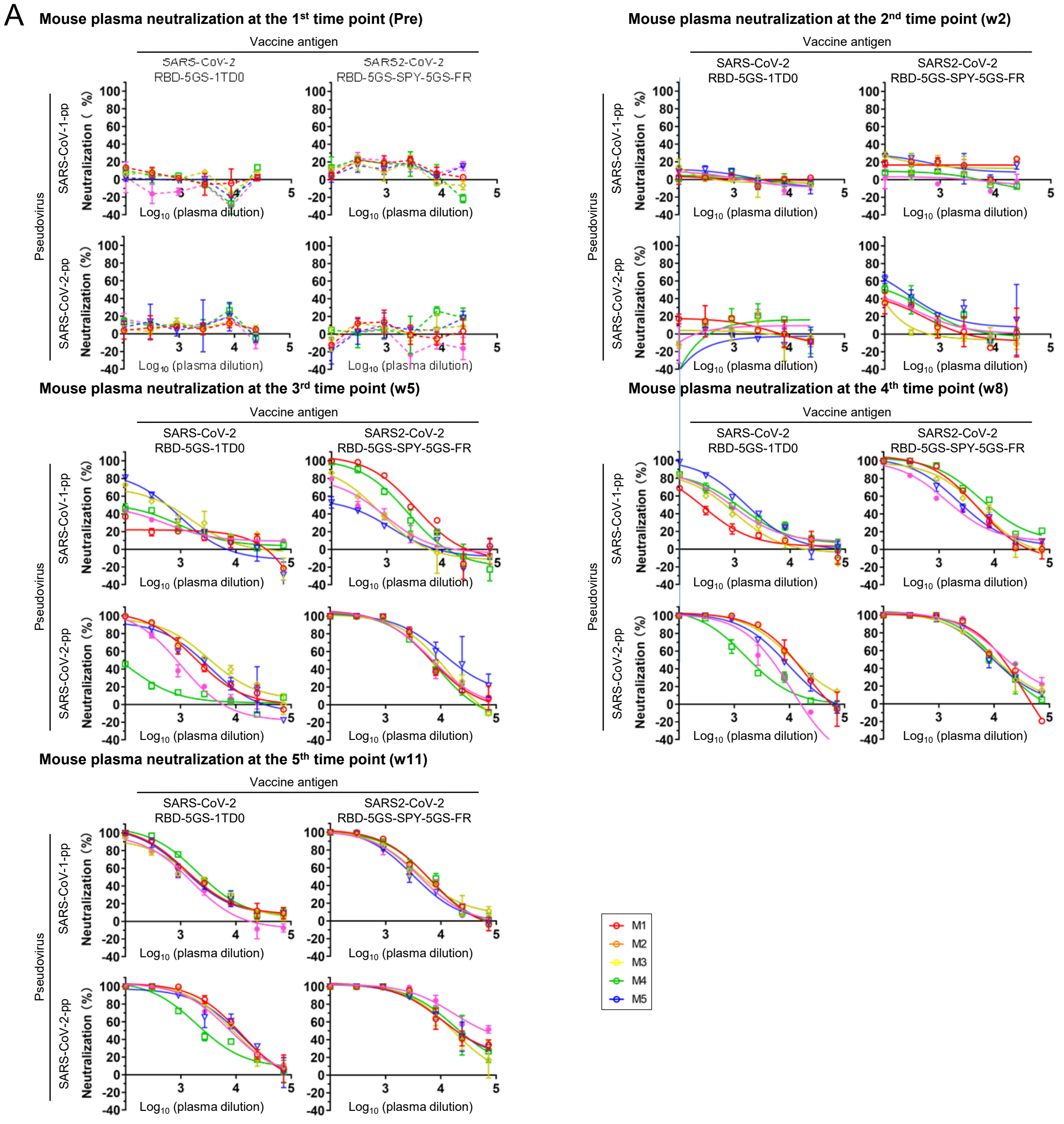


B **Mouse serum neutralization $ID_{50}$ titers**

Pseudovirus — SARS-CoV-1-pp

| SARS-CoV-2 vaccine antigen | W2 M1 | W2 M2 | W2 M3 | W2 M4 | W2 M5 | W5 M1 | W5 M2 | W5 M3 | W5 M4 | W5 M5 | W8 M1 | W8 M2 | W8 M3 | W8 M4 | W8 M5 | W11 M1 | W11 M2 | W11 M3 | W11 M4 | W11 M5 |
|---|---|---|---|---|---|---|---|---|---|---|---|---|---|---|---|---|---|---|---|---|
| RBD-5GS-1TD0 | <100 | <100 | <100 | <100 | <100 | 72 | 428 | 459 | 166 | 124 | 250 | 570 | 1608 | 1037 | 859 | 1979 | 1710 | 1816 | 2759 | 1183 |
| RBD-5GS-SPY-5GS-FR | <100 | <100 | <100 | <100 | <100 | 2870 | 482 | 189 | 1543 | 427 | 4456 | 4199 | 2858 | 8390 | 1941 | 5389 | 4986 | 3418 | 5401 | 4066 |

Pseudovirus — SARS-CoV-2-pp

| SARS-CoV-2 vaccine antigen | W2 M1 | W2 M2 | W2 M3 | W2 M4 | W2 M5 | W5 M1 | W5 M2 | W5 M3 | W5 M4 | W5 M5 | W8 M1 | W8 M2 | W8 M3 | W8 M4 | W8 M5 | W11 M1 | W11 M2 | W11 M3 | W11 M4 | W11 M5 |
|---|---|---|---|---|---|---|---|---|---|---|---|---|---|---|---|---|---|---|---|---|
| RBD-5GS-1TD0 | <100 | <100 | <100 | <100 | <100 | 2069 | 3655 | 2043 | <100 | 746 | 10882 | 12545 | 7382 | 2010 | 3245 | 10639 | 8742 | 9094 | 3047 | 8031 |
| RBD-5GS-SPY-5GS-FR | <100 | <100 | 220 | 143 | <100 | 6923 | 7926 | 16669 | 6079 | 7400 | 11048 | 11051 | 10012 | 10613 | 18178 | 20964 | 16955 | 20948 | 22920 | 51997 |

**fig. S7. SARS-CoV-2 RBD/RBD-SApNP vaccine-induced neutralizing antibody responses. (A)** Neutralization curves of mouse plasma from two SARS-CoV-2 RBD-based vaccine groups against SARS-CoV-1/2-pps. **(B)** Summary of $ID_{50}$ titers measured for two SARS-CoV-2 RBD-based vaccine groups against SARS-CoV-1/2-pps. Color coding indicates the level of $ID_{50}$ titer (white: no neutralization; green to red: low to high). The $ID_{50}$ values were calculated in GraphPad Prism 8.4.3, with the lower/upper constraints of %neutralization set at 0.0/100.0.

**fig. S8**

A

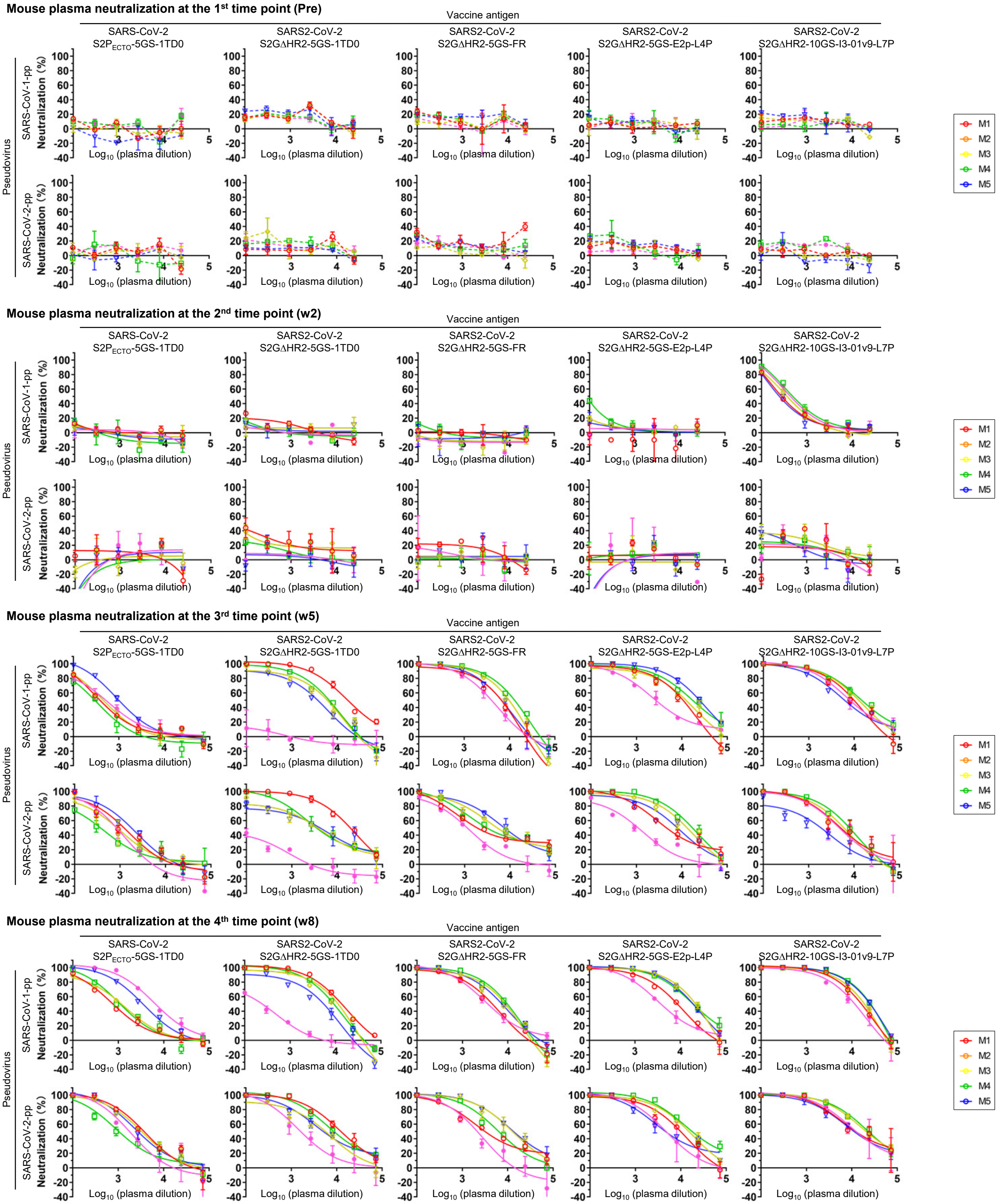

Mouse plasma neutralization at the 1st time point (Pre)
Mouse plasma neutralization at the 2nd time point (w2)
Mouse plasma neutralization at the 3rd time point (w5)
Mouse plasma neutralization at the 4th time point (w8)
Vaccine antigen
SARS-CoV-2 S2P$_{ECTO}$-5GS-1TD0
SARS2-CoV-2 S2GΔHR2-5GS-1TD0
SARS2-CoV-2 S2GΔHR2-5GS-FR
SARS2-CoV-2 S2GΔHR2-5GS-E2p-L4P
SARS2-CoV-2 S2GΔHR2-10GS-I3-01v9-L7P
Pseudovirus
SARS-CoV-1-pp
SARS-CoV-2-pp
Neutralization (%)
Log$_{10}$ (plasma dilution)
M1
M2
M3
M4
M5

### fig. S8

Mouse plasma neutralization at the 5th time point (w11)

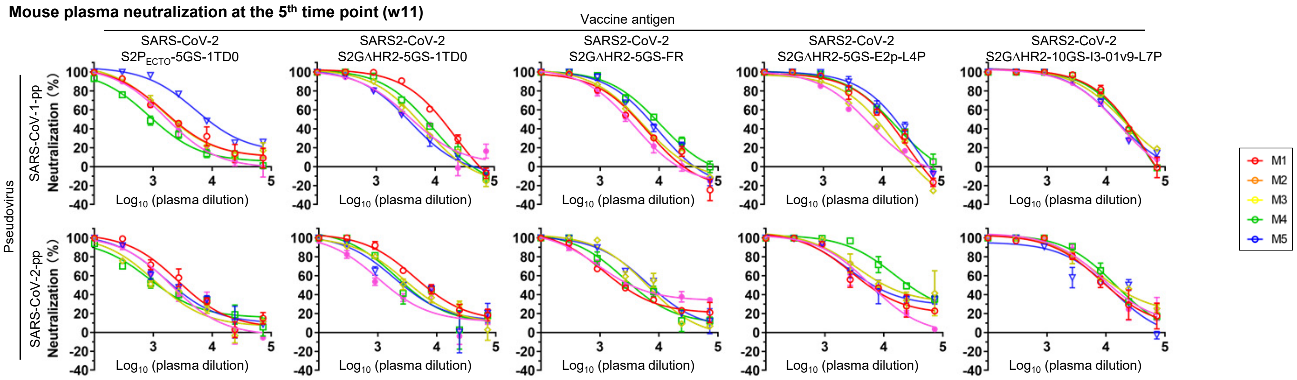


B **Mouse plasma neutralization $ID_{50}$ titers**

Pseudovirus: SARS-CoV-1-pp

| SARS-CoV-2 vaccine antigen | W2 | | | | | W5 | | | | | W8 | | | | | W11 | | | | |
|---|---|---|---|---|---|---|---|---|---|---|---|---|---|---|---|---|---|---|---|---|
| | M1 | M2 | M3 | M4 | M5 | M1 | M2 | M3 | M4 | M5 | M1 | M2 | M3 | M4 | M5 | M1 | M2 | M3 | M4 | M5 |
| $S2P_{ECTO}$-5GS-1TD0 | <100 | <100 | <100 | <100 | <100 | 405 | 444 | 1032 | 270 | 501 | 777 | 1062 | 3410 | 1121 | 6783 | 2588 | 2431 | 9374 | 1074 | 1806 |
| S2GDHR2-5GS-1TD0 | <100 | <100 | <100 | <100 | <100 | 17433 | 3960 | 2747 | 5170 | <100 | 13124 | 7579 | 3443 | 7902 | 221 | 11228 | 4377 | 3271 | 6009 | 4176 |
| S2GDHR2-5GS-FR | <100 | <100 | <100 | <100 | <100 | 4152 | 6773 | 5414 | 10007 | 3527 | 3735 | 5968 | 6443 | 7605 | 3872 | 3384 | 4103 | 5704 | 8194 | 2697 |
| S2GDHR2-5GS-E2p-L4P | <100 | <100 | <100 | <100 | <100 | 7954 | 10197 | 23509 | 17140 | 2915 | 7676 | 18213 | 17917 | 19538 | 4017 | 9170 | 5651 | 13334 | 12484 | 4799 |
| S2GDHR2-10GS-I3-01v9-L7P | 278 | 330 | 247 | 486 | 417 | 8250 | 11666 | 6695 | 15034 | 8694 | 14174 | 14888 | 22024 | 21474 | 9393 | 18257 | 18901 | 14175 | 17014 | 13137 |

Pseudovirus: SARS-CoV-2-pp

| SARS-CoV-2 vaccine antigen | w2 | | | | | w5 | | | | | w8 | | | | | w11 | | | | |
|---|---|---|---|---|---|---|---|---|---|---|---|---|---|---|---|---|---|---|---|---|
| | M1 | M2 | M3 | M4 | M5 | M1 | M2 | M3 | M4 | M5 | M1 | M2 | M3 | M4 | M5 | M1 | M2 | M3 | M4 | M5 |
| $S2P_{ECTO}$-5GS-1TD0 | <100 | <100 | <100 | <100 | <100 | 1050 | 787 | 1524 | 369 | 667 | 3515 | 3169 | 2897 | 1141 | 1683 | 3431 | 1428 | 2509 | 1494 | 1994 |
| S2GDHR2-5GS-1TD0 | <100 | <100 | <100 | <100 | <100 | 18419 | 3271 | 3238 | 4382 | <100 | 12641 | 4656 | 6148 | 9660 | 1707 | 7026 | 4096 | 3226 | 3491 | 1686 |
| S2GDHR2-5GS-FR | <100 | <100 | <100 | <100 | <100 | 3065 | 6659 | 8044 | 5164 | 1072 | 4156 | 12681 | 14238 | 6453 | 1827 | 3170 | 6292 | 7318 | 4051 | 5202 |
| S2GDHR2-5GS-E2p-L4P | <100 | <100 | <100 | <100 | <100 | 5778 | 11650 | 7370 | 16399 | 977 | 8613 | 14120 | 5999 | 19054 | 4733 | 6849 | 12699 | 8468 | 28041 | 6138 |
| S2GDHR2-10GS-I3-01v9-L7P | <100 | <100 | <100 | <100 | <100 | 5465 | 5151 | 1440 | 6831 | 4722 | 14684 | 22060 | 12830 | 23608 | 13575 | 10747 | 16081 | 8855 | 14951 | 12578 |

**fig. S8. SARS-CoV-2 spike/spike-SApNP vaccine-induced neutralizing antibody responses. (A)** Neutralization curves of mouse plasma from five SARS-CoV-2 spike-based vaccine groups against SARS-CoV-1/2-pps. **(B)** Summary of $ID_{50}$ titers measured for five SARS-CoV-2 spike-based vaccine groups against SARS-CoV-1/2-pps. Color coding indicates the level of $ID_{50}$ titer (white: no neutralization; green to red: low to high). The $ID_{50}$ values were calculated in GraphPad Prism 8.4.3, with the lower/upper constraints of %neutralization set at 0.0/100.0.

**fig. S9**

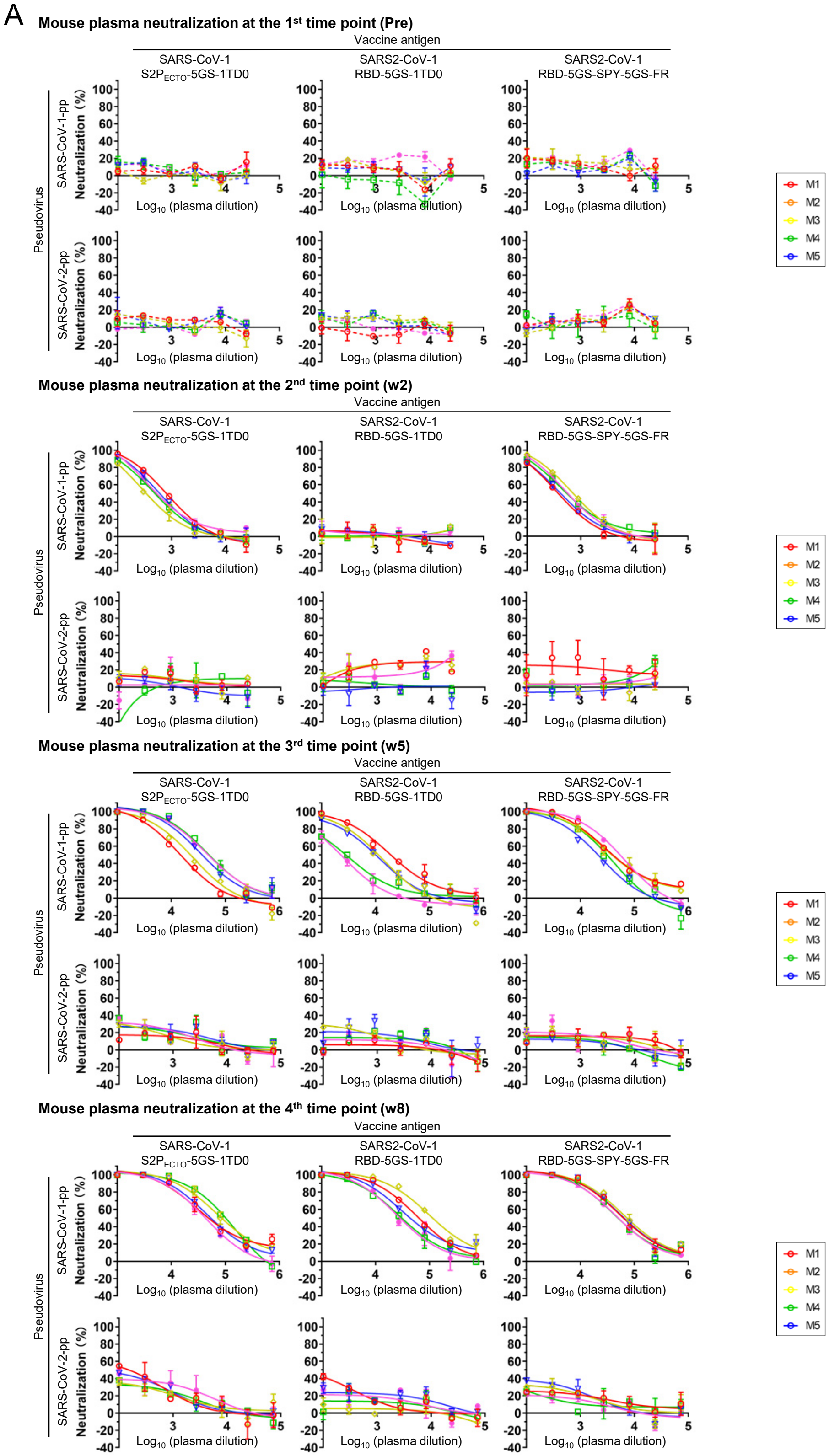
A
Mouse plasma neutralization at the 1st time point (Pre)
Mouse plasma neutralization at the 2nd time point (w2)
Mouse plasma neutralization at the 3rd time point (w5)
Mouse plasma neutralization at the 4th time point (w8)
Vaccine antigen
SARS-CoV-1 S2P$_{ECTO}$-5GS-1TD0
SARS2-CoV-1 RBD-5GS-1TD0
SARS2-CoV-1 RBD-5GS-SPY-5GS-FR
Pseudovirus
SARS-CoV-1-pp
SARS-CoV-2-pp
Neutralization (%)
Log$_{10}$ (plasma dilution)
M1
M2
M3
M4
M5

**Mouse plasma neutralization at the 5th time point (w11)**

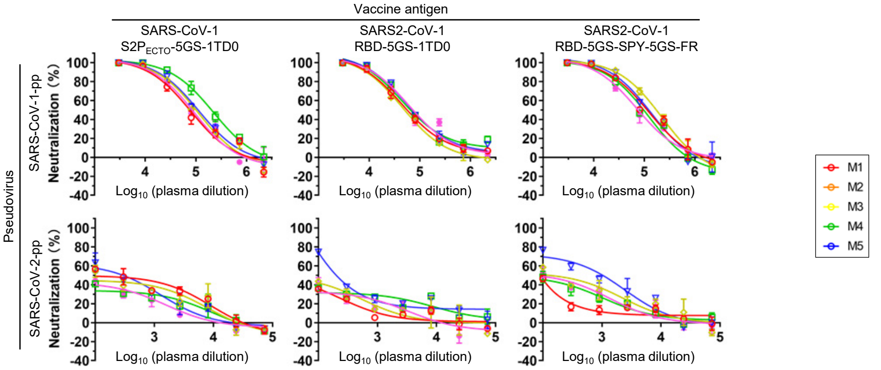


B **Mouse serum neutralization $ID_{50}$ titers**

Pseudovirus: SARS-CoV-1-pp

| SARS-CoV-1 vaccine antigen | w2 | | | | | w5 | | | | | w8 | | | | | w11 | | | | |
|---|---|---|---|---|---|---|---|---|---|---|---|---|---|---|---|---|---|---|---|---|
| | M1 | M2 | M3 | M4 | M5 | M1 | M2 | M3 | M4 | M5 | M1 | M2 | M3 | M4 | M5 | M1 | M2 | M3 | M4 | M5 |
| S2P$_{ECTO}$-5GS-1TD0 | 770 | 343 | 591 | 487 | 593 | 14583 | 21202 | 38507 | 51532 | 49913 | 56161 | 91290 | 57692 | 90786 | 42757 | 74328 | 94977 | 109706 | 198568 | 80619 |
| RBD-5GS-1TD0 | <100 | <100 | <100 | <100 | <100 | 19980 | 10126 | 9368 | 2830 | 2110 | 65863 | 120732 | 49566 | 32475 | 30098 | 62357 | 52065 | 78969 | 74719 | 80296 |
| RBD-5GS-SPY-5GS-FR | 370 | 719 | 423 | 588 | 584 | 45123 | 43109 | 19859 | 26996 | 50574 | 66993 | 77427 | 69400 | 53156 | 52677 | 109989 | 157931 | 117525 | 86514 | 77985 |

Pseudovirus: SARS-CoV-2-pp

| SARS-CoV-1 vaccine antigen | w2 | | | | | w5 | | | | | w8 | | | | | w11 | | | | |
|---|---|---|---|---|---|---|---|---|---|---|---|---|---|---|---|---|---|---|---|---|
| | M1 | M2 | M3 | M4 | M5 | M1 | M2 | M3 | M4 | M5 | M1 | M2 | M3 | M4 | M5 | M1 | M2 | M3 | M4 | M5 |
| S2P$_{ECTO}$-5GS-1TD0 | <100 | <100 | <100 | <100 | <100 | <100 | <100 | <100 | <100 | <100 | 174 | <100 | 142 | <100 | 140 | 325 | 216 | 268 | 112 | 114 |
| RBD-5GS-1TD0 | <100 | <100 | <100 | <100 | <100 | <100 | <100 | <100 | <100 | <100 | <100 | <100 | <100 | <100 | <100 | <100 | 103 | 272 | <100 | <100 |
| RBD-5GS-SPY-5GS-FR | <100 | <100 | <100 | <100 | <100 | <100 | <100 | <100 | <100 | <100 | <100 | <100 | <100 | <100 | <100 | <100 | 240 | 717 | 150 | 184 |

C

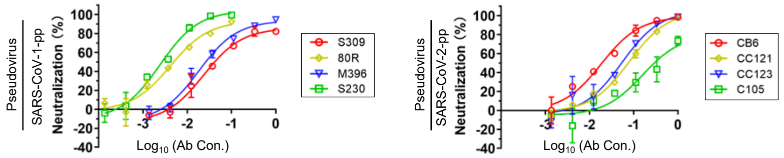


D **Mouse serum neutralization against MLV-pp at the 4th time point (w8)**

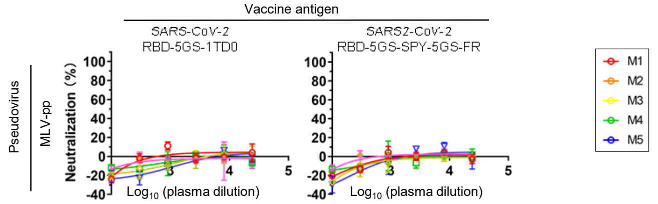


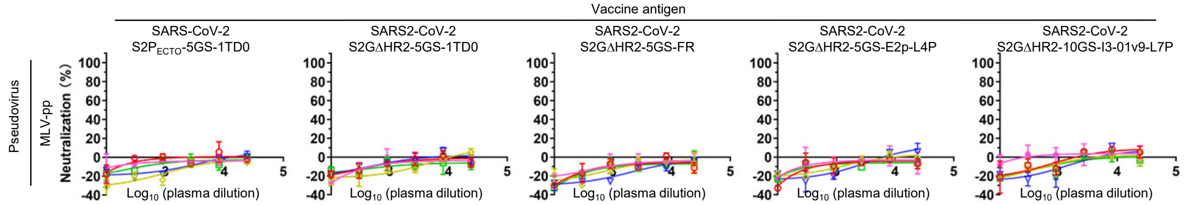


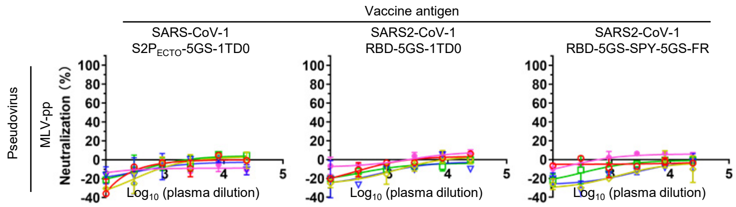

**fig. S9. SARS-CoV-1 spike/RBD/RBD-SApNP vaccine-induced neutralizing antibody responses. (A)** Neutralization curves of mouse plasma from three SARS-CoV-1 vaccine groups against two SARS-CoV-1/2-pps. **(B)** Summary of $ID_{50}$ titers measured for three SARS-CoV-1 vaccine groups against SARS-CoV-1/2-pps. Color coding indicates the level of $ID_{50}$ titer (white: no neutralization; green to red: low to high). The $ID_{50}$ values were calculated in GraphPad Prism 8.4.3, with the lower/upper constraints of %neutralization set at 0.0/100.0. **(C)** Neutralization curves of four known SARS-CoV-1/2 NAbs against respective pseudoviruses. SARS-CoV-1-pp (left) and SARS-CoV-2-pp (right). **(D)** Neutralization curves of mouse plasma from all vaccine groups at w8 against MLV-pps as a negative control. No MLV-pp neutralization was observed.

### Table S1

**Table S1. Key *in vitro* and *in vivo* characteristics of RBD and spike-based SARS-CoV-2 vaccines [a]**

| | RBD-based constructs | | | | Spike-based constructs | | | | |
|---|---|---|---|---|---|---|---|---|---|
| | RBD-5GS-1TD0 | RBD-5GS-SPY-5GS-FR | RBD-5GS-SPY-5GS-E2p-LD4-PADRE | RBD-5GS-SPY-5GS-I3-01v9-LD7-PADRE | $S2P_{ECTO}$-5GS-1TD0 | S2GΔHR2-5GS-1TD0 | S2GΔHR2-5GS-FR | S2GΔHR2-5GS-E2p-LD4-PADRE | S2GΔHR2-10GS-I3-01v9-LD7-PADRE |
| Display scaffold [b] | 1TD0 | Ferritin | E2p | I3-01v9 | 1TD0 | 1TD0 | Ferritin | E2p | I3-01v9 |
| Multilayered (Y/N) [c] | NA | N | Y | Y | NA | NA | N | Y | Y |
| PADRE included (Y/N) [d] | NA | N | Y | Y | NA | NA | N | Y | Y |
| Antigen (Ag) type [e] | Monomer | Monomer | Monomer | Monomer | Trimer | Trimer | Trimer | Trimer | Trimer |
| Number of Ags [f] | 3 | 24 | 60 | 60 | 1 | 1 | 8 | 20 | 20 |
| Display method [g] | Fusion | SPY | SPY | SPY | Fusion | Fusion | Fusion | Fusion | Fusion |
| Number of plasmids [h] | 1 | 2 | 2 | 2 | 1 | 1 | 1 | 1 | 1 |
| Protein yield [i] | +++++ | ++++ | + | +++ | +++ | +++ | +++ | ++ | +++ |
| Protein purity [j] | +++++ | ++++ | + | +++ | ++++ | +++++ | +++ | ++ | +++ |
| ELISA binding [k] | ++++ | +++++ | ND | ND | ++++ | ++++ | +++++ | +++++ | +++++ |
| BLI binding [l] | ++ | +++++ | ND | ND | ++ | ++ | +++ | +++++ | +++++ |
| W8 binding Ab titer [m] | ++++ | +++++ | ND | ND | ++ | +++ | +++ | ++++ | +++++ |
| W8 NAb titer [n] | ++ | +++ | ND | ND | ++ | +++ | +++ | ++++ | +++++ |
| T-cell response [o] | ND | ND | ND | ND | ++ | ND | ND | +++ | ++++ |

[a] Summary of key vaccine attributes with "+" indicating positive outcome (+ being broadline and +++++ being optimal), "NA" not applicable, and "ND" not determined.
[b] Display scaffold – A trimerization motif (1TD0) and three self-assembling protein nanoparticles (SApNPs), ferritin, E2p, and I3-01, are used to display the RBD and spike.
[c] Multilayered (Y/N) – Whether the SApNP construct contains an inner layer of locking domains (LD) and a cluster of T-cell epitopes (PADRE). Y: yes, N: no.
[d] PADRE included (Y/N) – Whether the SApNP construct contains a pan DR epitope that activates $CD4^+$ T cells.
[e] Antigen (Ag) type – The RBD is a monomer, whereas the spike is a trimer of cleaved S1/S2 heterodimers or uncleaved S proteins.
[f] Number of Ags – Number of antigens in the properly folded and assembled vaccine immunogen.
[g] Display method – The antigen is either conjugated to SApNPs using the SpyTag/SpyCatcher (SPY) system or genetically fused to a trimerization motif or SApNP subunit.
[h] Number of plasmids – The number of expression plasmids needed to produce the vaccine immunogen in ExpiCHO cells.
[i] Protein yield – Protein yield is measured after purification on a CR3022 antibody column. Of note, individual antigens usually have higher yield than their SApNPs, and the 24-meric SApNP (ferritin) tends have greater yield than the 60-meric SApNPs, E2p and I3-01.
[j] Protein purity – Protein purity is measured by qualitatively comparing the ratio of various species (aggregate, SApNP, trimer, dimer and monomer) in the SEC profile.
[k] ELISA binding – $EC_{50}$ values are calculated from ELISA binding to a panel of known antibodies.
[l] BLI binding – Binding signals (nm) are compared as most constructs show negligible off-rates in the Octet experiments.
[m] W8 binding Ab titer – $EC_{50}$ titers are compared at week 8, where serum binding antibody response reaches the plateau.
[n] W8 NAb titer – $ID_{50}$ titers are compared at week 8, where serum NAb response reaches the plateau.
[o] T-cell response – End-point T-cell (Th1, cytolytic $CD4^+$, and CD8) responses.
